# Algal Exudates Promote Conjugation in Marine Roseobacters

**DOI:** 10.1101/2023.11.14.567099

**Authors:** Yemima Duchin Rapp, Valeria Lipsman, Lilach Yuda, Ilya V Kublanov, Dor Matsliyah, Einat Segev

## Abstract

Horizontal gene transfer (HGT) is a pivotal mechanism driving bacterial evolution, conferring adaptability within dynamic marine ecosystems. Among HGT mechanisms, conjugation mediated by type IV secretion systems (T4SSs) plays a central role in the ecological success of marine bacteria. However, the conditions promoting conjugation events in the marine environment are not well understood. Roseobacters, abundant marine bacteria commonly associated with algae, possess a multitude of T4SSs. Many Roseobacters are heterotrophic bacteria that rely on algal secreted compounds to support their growth. These compounds attract bacteria, facilitating colonization and attachment to algal cells. Algae and their metabolites bring bacteria into close proximity, potentially promoting bacterial HGT. Investigation across various Roseobacters revealed that algal exudates indeed enhance plasmid transfer through conjugation. While algal exudates do not influence the transcription of bacterial conjugative machinery genes, they promote bacterial attachment, potentially stabilizing proximity and facilitating HGT. Notably, under conditions where attachment is less advantageous, the impact of algal exudates on conjugation is reduced. These findings suggest that algae enhance bacterial conjugation primarily by fostering attachment and highlight the importance of studying bacterial HGT within the context of algal-bacterial interactions.

## Introduction

Horizontal gene transfer (HGT), a key process in bacterial evolution, is the transfer of genetic material between a donor to an unrelated recipient. Bacteria, while maintaining small size genomes^1^, display extraordinary genetic variations that enable them extreme flexibility and adaptability^1,2^. The ability of bacteria to obtain fragments of DNA in one single HGT event supports this flexibility^2^. HGT allows bacteria to adapt to environmental changes by acquiring new metabolic capabilities that facilitate various abilities such as symbiosis^3^, antibiotic resistance^4^, metabolic activity^5^ and pathogenicity^6^ thereby enabling bacteria to inhabit new and dynamic niches.

Horizontal transfer of genetic material is achieved by several main mechanisms, including 1) Transformation, which is the uptake of extracellular DNA^7^, 2) Transduction, which is bacteriophage-mediated transfer^7^, 3) Gene transfer agents (GTAs), that are bacteriophage-like particles that carry random pieces of the producing bacterial genome^8^, and 4) Conjugation, which is the transfer of a plasmid between bacterial cells through direct cell-to-cell contact^7–9^. Conjugation typically occurs through the type IV secretion system (T4SS). T4SS is a versatile system found in both Gram-negative and Gram-positive bacteria. The T4SS enables the delivery of proteins and DNA between different bacterial strains, species, and even from bacteria to eukaryotic cells^10^. It is involved in a verity of processes such as toxin delivery, DNA release, DNA uptake, effector molecule translocation and conjugation^11^. The T4SS also facilitates cell-to-cell contact and biofilm formation between bacteria. In turn, cell-to-cell contact and a sessile lifestyle in a biofilm can improve the efficiency of the conjugative transfer^12–20^.

HGT, and specifically conjugation that was demonstrated between Roseobacters^6,21–23^, has been suggested as a central reason for the ecological success of Roseobacters^22^. These bacteria exhibit diverse metabolic capabilities and a wide range of symbiotic lifestyles^24^. Many Roseobacters engage in various interactions with algal hosts^25–30^. Algae exude various metabolites that support bacterial growth in the nutrient-poor oligotrophic ocean^24,25,31^. Thus, marine bacteria can secure a nutrient flux through proximity to an algal host. Algal exudates were shown to influence bacterial physiology^24–26,31,32^ and shape algal-bacterial interactions^25,33^. In the vicinity of algal hosts, bacteria experience spatial proximity to one another, with certain bacteria attaching to the surface of the algal cells^25,32^. Since bacterial proximity, cell-to-cell contact, and attachment can promote HGT^13,21,34,35^, algal exudates may promote attachment and consequently bacterial HGT.

Here, we sought to explore Roseobacter conjugation in the ecological context of an algal host. Therefore, our study employed Roseobacter species that were identified in environmental samples of algae^25,36^. As a model algal host, we used the unicellular alga *Emiliania huxleyi* (also termed G*ephyrocapsa huxleyi*)^37^, which is widespread in modern oceans and was shown to harbor a rich community of bacteria, including Roseobacters^25,36,38^. First, we tested the ability of various Roseobacters to successfully conduct conjugation. Next, we examined the impact of algal exudates on bacterial conjugation. To understand the underlying mechanism, we monitored the expression of T4SS-encoding genes and bacterial attachment capabilities following exposure to algal exudates. Our findings demonstrate improved bacterial conjugation efficiency following exposure to algal exudates. While T4SS gene expression did not increase in response to algal exudates, bacterial attachment capabilities did improve. Conducting conjugation assays under conditions in which attachment is not beneficial, resulted in a reduced impact of the algal exudates. Our data suggest that algal exudates enhance bacterial conjugation primarily by promoting bacterial attachment.

## Materials & Methods

### Bacterial strains and growth conditions

The bacterial strains *Phaeobacter inhibens* DSM 17395 and *Dinoroseobacter shibae* DFL-12 were purchased from the German Collection of Microorganism and Cell Cultures (DSMZ, Braunschweig, Germany) (Table S1). *P. inhibens* P72 was kindly provided by Prof. Juan Barja from University of Santiago de Compostela, Spain, and Dr. Jӧrn Petersen from the Leibniz Institute, DSMZ, Germany (Table S1). The bacterial strains of *Phaeobacter inhibens* DSM 17395 with a kanamycin-resistance cassette on its chromosome (*P. inhibens* DSM 17395^chr-kana^), *Marinovum algicola* DG898 DSM 27768 with a gentamicin-resistance cassette on its chromosome (*M*. *algicola* DG898 ^chr-gent^), *Dinoroseobacter shibae* DFL-12 with a gentamicin-resistance cassette on its native 126 kb plasmid (*D. shibae* DFL-12 ^p126-gent^), *Dinoroseobacter shibae* DFL-12 with a gentamicin-resistance cassette on its chromosome (*D. shibae* DFL-12 ^chr-gent^), *P. inhibens* P72 with a kanamycin-resistance cassette on its native 57 kb plasmid (*P. inhibens* P72 ^p57-kana^), and *P. inhibens* P72 with a kanamycin-resistance cassette on its chromosome (*P. inhibens* P72 ^chr-kana^) were kindly provided by Dr. Jӧrn Petersen from the Leibniz Institute, DSMZ, Germany (Table. S1).

*D. shibae* cultures were plated on ½ marine broth 2216 (MB) (Difco, USA) agar plates (MB, 18.7 g/L; agar, 16 g/L) with 30 µg/ml gentamicin antibiotic. *M. algicola* cultures were plated on ½ YTSS agar plates (yeast extract, 2 g/L; trypton, 1.25 g/L; sea salts, 20 g/L; agar, 16 g/L, all purchased from Sigma-Aldrich) with 30 µg/ml gentamicin antibiotic. *P. inhibens* DSM 17395 cultures were plated on ½ YTSS agar plates with 30 µg/ml gentamicin or 150 µg/ml kanamycin antibiotic. *P. inhibens* P72 was plated on ½ YTSS agar plates with 150 µg/ml kanamycin antibiotic. Transconjugants of *D. shibae* were plated on ½ MB agar plates with 30 µg/ml gentamicin and 150 µg/ml kanamycin antibiotics. Transconjugants of *P. inhibens* DSM 17395, *P. inhibens* P72 and *M. algicola* were plated on ½ YTSS agar plates with 30µg/ml gentamicin and 150 µg/ml kanamycin antibiotics.

Cultures of algae and bacteria were grown in artificial seawater (ASW) prepared according to Goyet and Poisson^39^. ASW contained mineral salts (NaCl, 409.41 mM; Na_2_SO_4_, 28.22 mM; KCl, 9.08 mM; KBr, 0.82 mM; NaF, 0.07 mM; Na_2_CO_3_, 0.20 mM; NaHCO_3_, 2 mM; MgCl_2_, 50.66 mM; SrCl_2_, 0.09 mM), L1 trace elements (FeCl_3_ · 6H_2_O, 3.15 mg/L; Na_2_EDTA · 2H_2_O, 4.36 mg/L; CuSO_4_ · 5H2O, 9.8 μg/L; Na_2_MoO_4_ · 2H_2_O, 6.3 μg/L; ZnSO_4_ · 7H_2_O, 22 μg/L; CoCl_2_ · 6H_2_O, 10 μg/L; MnCl_2_ · 4H_2_O, 180 μg/L) and L1 nutrients (NaNO_3_, 882 μM; NaH_2_PO_4_, 36.22 μM). The components were dissolved in Milli-Q water (IQ 7003; Merck, Darmstadt, Germany) and the pH adjusted to 8 with HCl. Stock solutions of L1 trace elements and L1 nutrients were purchased from the Bigelow Laboratory for Ocean Sciences (Boothbay, ME, USA).

Cultures of *P. inhibens* DSM 17395, *P. inhibens* P72, and *M. algicola* were grown in ASW medium supplemented with additional carbon (glucose, 5.5 mM, Sigma Aldrich), nitrogen (NH_4_Cl, 5 mM, Sigma Aldrich), sulfur (Na_2_SO_4_, 33 mM, Merck), and f/2 vitamins (thiamine HCl, 100 μg/L; biotin, 0.5 μg/L; vitamin B_12_, 0.5 μg/L, Bigelow Laboratory for Ocean Sciences). *D. shibae* cultures were cultivated with minor modifications including 5.5 mM succinate as a carbon source (Sigma Aldrich) and different vitamins mix (adjusted from Gonzalez *et al.,*^40^ Thiamine HCl, 0.148 μM/L; biotin, 0.082 μM/L; Pyridoxine HCL, 0.486 μM/L; Nicotinic acid, 0.406 μM/L; Pantothenic acid, 0.228 μM/L; Cyanocobalamin, 0.0006 μM/L; pABA, 0.364 μM/L). This medium, ASW supplemented with vitamins and nutrients, is referred to as CNS. Bacteria were cultivated at 30 °C under constant shaking at 130 rpm.

### Algal strain, growth conditions and monitoring

The axenic algal strain of *Emiliania huxleyi* CCMP3266 (also termed G*ephyrocapsa huxleyi*^37^) was purchased from the National Center for Marine Algae and Microbiota (Bigelow Laboratory for Ocean Sciences, Maine, USA). Algae were grown in ASW (as described above) supplemented with f/2 vitamins. Algae were grown in standing cultures in a growth room at 18°C under a light/dark cycle of 16/8 h. Illumination intensity during the light period was 150 mmoles/m^2^/s. Absence of bacteria in axenic algal cultures was monitored periodically both by plating on ½ YTSS plates and under the microscope.

Cultures of *E. huxleyi* were grown as follows; algal cells concentrations from a late exponential phase culture were counted by a CellStream CS-100496 flow cytometer (Merck, Darmstadt, Germany) using 561 nm excitation and 702 nm emission. Algal cells were gated according to event size and fluorescence intensity. An inoculum of 10^5^ algal cells was introduced into 100 ml of ASW and incubated as described above. Algal growth in cultures was monitored by a CellStream flow cytometer as described. For each biological replicate 50,000 events were recorded.

### Algal filtrates

To obtain algal filtrates, algal cultures were grown as described above for 12 days (Fig. S6). An initial washing of the filters was conducted by passing 100 ml ASW medium through the filter (Thermo scientific, 0.22 μm PES, 250ml). Then, algal cultures were vacuum filtered, and filtrates pH was adjusted to pH 8 (Thermo Scientific Eutech pH 700 Meter) with HCl, for identical pH levels across both control and treatment. This step was followed by a second filtration since the titration procedure was conducted in non-sterile conditions. The ASW medium used as control was subjected to the same procedure. To validate that vacuum filtration did not rupture the algal cells, the filters were suspended in ASW and the algal population was assessed using flow cytometry.

### Monitoring bacterial growth

Bacterial cultures were monitored daily by OD_600_ measurements using plastic cuvettes in a spectrophotometer (Ultrospec 2100 pro, Biochrom).

### Conjugation assay

*D. shibae* DFL-12 ^p126-gent^, *P. inhibens* P72 ^p57-kana^*, P. inhibens* DSM 17395 ^p262-gent^, *P. inhibens* DSM 17395 ^p78-gent^ and *P. inhibens* DSM 17395 ^p65-gent^ with plasmids carrying antibiotic resistance genes were used as donor strains for conjugation. *P. inhibens* DSM 17395^chr-gent^, *P. inhibens* DSM 17395^chr-kana^*, P. inhibens* P72 ^chr-kana^, *M*. *algicola* DG898 ^chr-gent^, and *D. shibae* DFL-12 ^chr-gent^ with chromosomes carrying antibiotic resistance genes were used as recipient strains for conjugation (Table S2). The conjugation assay involved several stages: first, preparing pre-cultures of pure donors and recipients; then, mating in liquid or agar plates; and finally selection on selective agar plates.

#### Pre-cultures

Bacteria were inoculated from a frozen stock (-80 °C) onto selective agar plates and incubated 2-4 days at 30 °C. Then, bacteria were transferred from the plates to ASW medium and adjusted to an OD_600_ of 0.1. Next, bacteria were diluted to an OD_600_ of 0.001 in the medium previously specified for each strain. Pre-cultures were grown to an OD_600_ of 0.3, followed by centrifugation of 10 min at 3,200 g. Bacteria were washed with ASW and centrifuged again for 10 min at 3,200 g. Samples were adjusted to an OD_600_ of 25 using ASW.

#### Mating

For conjugation assay in liquid media (Fig. 1, Fig. 2), the treatment consisted of 75% fresh algal filtrates diluted in 25% ASW (v/v, ratio of 3:1). Both control (ASW) and treated cultures were supplemented with L1 trace elements, L1 nutrients, 5.5 mM glucose, vitamin mix, 33 mM Na_2_SO_4_, and 5 mM NH_4_Cl. The final control and treatment medium are referred to as CNS, with the treatment containing 75% volume of algal filtrates and 25% volume of ASW (v/v), in contrast to the control which consists solely of ASW. To each 14 ml tube (Corning Falcon Polypropylene round bottom tube), 2.4 ml of the medium, 100 μl of the donor strain, and 20 μl of the recipient strain were added. The tubes were mixed and incubated at 30 °C for 24 h without shaking.

For mating on agar plates (Fig. 6), agar (16 g) was added to 250 ml of ASW medium (w/v) and autoclaved. After cooling, 750 ml of algal filtrates or ASW were added as treatment or control respectively (v/v). Both control and treated cultures were supplemented with L1 trace elements, L1 nutrients, vitamin mix, Glucose, Na_2_SO_4_, and NH_4_Cl (consisting of the CNS medium). The mediums were mixed and poured onto the plates. Then, 20 μl of donor cells and 4 μl of recipient cells were mixed and plated on the agar plates, followed by 24 h incubation at 30 °C. For mating on MB agar plates (Fig. S3), 20 μl of donor cells and 4 μl of recipient cells were mixed and plated on MB agar plates, followed by 24 h incubation at 30 °C.

#### Selection and quantification

for conjugation in liquid media, each tube was plated in serial dilutions in two technical replicates on three selective plates: one for the donor, one for the recipient and one for the transconjugants. For conjugation on agar plates, bacteria were scraped from the plate, resuspended in 150 μl of ASW, and plated in serial dilutions as described for the conjugation assay in liquid. Plates were incubated for 48-96 h at 30 °C. Colonies were counted and quantified. Importantly, transconjugants can grow on selective plates containing a single antibiotic, used for quantifying the donors and recipients, as they possess resistance to both antibiotics. However, our data indicated that there were 2-5 orders of magnitude fewer transconjugants compared to donors and recipients. Therefore, the growth of transconjugants on single-antibiotic plates is negligible in comparison to cell numbers of donors or recipients.

#### Validation

From each donor-recipient conjugation pair, seven transconjugant colonies were re-streaked on double-antibiotics plates. After three rounds of re-streaking, single transconjugant colonies were subjected to DNA extraction using Wizard Genomic DNA Purification Kit (Promega), and the DNA was used for PCR amplification using the PCRBIO HS Taq Mix Red polymerase. For each transconjugant, three sets of primers were designed for validation: a primer set specific for the donor, a primer set specific for the recipient and primer set specific for the transferred plasmid (Table S3, Table S4).

### Quantitative Real Time-PCR (qRT-PCR)

Pre-cultures of *P. inhibens* P72 ^p57-kana^ and *D. shibae* DFL-12 ^p126-gent^ bacteria were grown as previously described. At OD_600_ of 0.3, cells were harvested by centrifugation at 3,200 g for 10 min at room-temperature and transferred into ASW (control) and algal filtrates (treatment). The treatment consisted of 75% fresh algal filtrates diluted in 25% of ASW (v/v). Both control and treated cultures were supplemented with L1 trace elements, L1 nutrients, vitamin mix, 33 mM Na_2_SO_4_, and 5 mM NH_4_Cl. The final control and treatment medium is referred to as CNS, with the treatment containing 75% volume of algal filtrates and 25% volume of ASW (v/v), in contrast to the control which consists solely of ASW. *P. inhibens* P72 ^p57-kana^ cultures were additionally supplemented with 5.5 mM Glucose, and *D. shibae* DFL-12 ^p126-gent^ cultures were supplemented with 5.5 mM succinate. After 30 min of incubation at 30 °C with shaking at 130 rpm, the cells were centrifuged at 3,200 g for 10 min at 4 °C. Cells were transferred into 450 µL of RLT buffer containing 1% β-mercapto-ethanol and kept at -80 °C until RNA extraction. RNA was extracted using the Isolate II RNA mini kit (Meridian Bioscience, London, UK) according to the manufacturer instructions. Cells were ruptured by adding low Binding Silica Grinding Beads (100 µm) (SPEX, Metuchen, Netherland) to the thawed sample and were then beaten for 5 min at 30 mHz. For further removal of genomic DNA, RNA samples were treated with 2 µL Turbo DNAse (ThermoFisher) in a 60 µL reaction volume. RNA samples were cleaned and concentrated using RNA Clean & Concentrator TM-5 kit (Zymo Research, Irvine, CA, USA) according to the manufacturer instructions. Equal concentrations of RNA were utilized for cDNA synthesis using Superscript IV (ThermoFisher), according to manufacturer instructions. qRT-PCR was conducted in 384-well plates using the SensiFAST SYBR Lo-ROX Kit (Meridian Bioscience) in a QuantStudio 5 (384-well plate) qRT-PCR cycler (Applied Biosystems, Foster City, CA, USA). The qRT-PCR program ran according to enzyme requirements for 40 cycles. Primers are listed in table S5. Primer efficiencies were calculated through standard curves using the QuantStudio 5 software. Gene expression ratios were analyzed as previously shown^41^, by geometric averaging of the housekeeping genes *gyrA*, and *recA*. To validate the absence of genomic DNA contamination, the same procedure and program were applied on RNA samples that were not reverse transcribed.

### Reporter strains

The T4SS reporter strains were constructed by cloning the *virB* promoter upstream of a gene encoding the super-folded green fluorescent protein (sfGFP) in the pBBR1MCS-5 vector^42^ (kindly provided by Prof. Kenneth M. Peterson, Louisiana State University Medical Center, USA) (Table 6).

#### Construction of *P. inhibens* P72 reporter strains

The *virB* promoter from the 184 kb plasmid was amplified using primers 987 and 988 (Table S4). The *virB* promoter from the 57 kb plasmid was amplified using primers 989 and 990 (Table S4). The amplified fragments were cloned separately into the pYDR1 vector (Table S6) upstream of a sfGFP encoding gene, using restriction-free cloning^43,44^. The resulting plasmids, pYDR3 for the 184 kb plasmid and pYDR10 for the 57 kb plasmid (Table S6) were introduced into competent *P. inhibens* P72 cells by electroporation. Preparations of competent cells was performed as previously described^45^, with slight modifications. Shortly, cells were grown to an OD_600_ of 0.7 in ½ YTS with 40 g/L of sea salts (also termed full salts medium). Bacteria were washed 3 times using 10% (v/v) ice-cold glycerol, centrifuged each time for 5 min at 4 °C and 5,500 g. The competent cells were subsequently adjusted to an OD_600_ of 20 using 10% (v/v) ice-cold glycerol, frozen in liquid nitrogen and stored at -80°C. Electroporation was conducted with 300 μl aliquot of electrocompetent cells, using 10 µg of DNA in a 2 mm cuvette at 2.5 V, followed by 4 h recovery in ½ YTS with full salts. The transformed cells were then plated on ½ YTSS plates containing 50 μg/ml gentamicin. Resistant colonies were re-streaked three times. Sequencing and PCR validation of the reporter strains was conducted using primers 132 and 255 (Table S5).

#### Construction of *D. shibae* reporter strains

The *virB* promoter from the 191 kb plasmid was amplified using primers 974 and 975 (Table S4). The *virB* promoter from the 126 kb plasmid was amplified using primers 972 and 973 (Table S4). The amplified fragments were separately cloned into pYDR1 vector (Table S6) upstream of a gene encoding the sfGFP protein, using restriction-free cloning^43,44^. The resulting plasmids, pYDR6 for the 191 kb plasmid and pYDR7 for the 126 kb plasmid (Table S6) were conjugated into *D. shiba*e as previously described^45^, with slight modifications. Shortly, the constructs were introduced into competent *E. coli* ST18 DSM 22074, carrying a *hemA* deletion that results in defective tetrapyrrole biosynthesis (purchased from DSMZ). Hence, the mutant strain requires addition of 5-aminolevulinic acid to grow. The donor *E. coli* ST18 containing the pYDR6 or pYDR7 plasmids was mated with the recipient *D. shibae.* Cultures of *D. shibae* recipients were cultivated for 24 h at 30 °C to an OD_600_ of 0.4 in MB medium. The donor *E. coli* ST18 was grown in LB medium supplemented with 50 μg/ml 5-aminolevulinic acid (Sigma-Aldrich) at 37 °C to an OD_600_ of 0.3. Both cultures were centrifuged, concentrated 25-fold, and mixed in a donor-to-recipient ratio of 10:1. Then, 50 μl drops of the mixed cultures were plated on ½ MB plates containing 50 μg/ml 5-aminolevulinic acid, and incubated for 24 h at 30 °C. Cells were scraped from the plate, resuspend in 100 μl of MB, and plated on ½ MB plates containing 80 μg/ml gentamicin. Colonies of transconjugants were re-streaked three times. Sequencing and PCR validation of the reporter strains was conducted using primers 132 and 255 (Table S5).

#### Reporter strain assay

Cultures of the reporter strains were adjusted to an OD_600_ of 0.001 and grown in ASW (control) and algal filtrates (treatment). The treatment consisted of 75% fresh algal filtrates diluted in 25% of ASW (v/v). Both control and treatment cultures were supplemented with L1 trace elements, L1 nutrients, vitamin mix, 33 mM Na_2_SO_4_, and 5 mM NH_4_Cl. *P. inhibens* P72 cultures were additionally supplemented with 5.5 mM Glucose, and *D. shibae* cultures were supplemented with 5.5 mM succinate. The final control and treatment medium is referred to as CNS, with the treatment containing 75% volume of algal filtrates and 25% volume of ASW (v/v), in contrast to the control which consists solely of ASW. Cultures were grown at 30 °C with shaking at 130 rpm and harvested at an OD_600_ of 0.3 for GFP visualization. Bacterial populations were monitored using a Merck CellStream flow cytometer, excited at 561 nm, collected at 615–789 nm. GFP was evaluated by determining a GFP fluorescence threshold (according to autofluorescence of control cultures) and calculating the percentage of bacterial cells with fluorescence intensity higher than the threshold. For each biological replicate 100,000 events were recorded. Results are presented as percentages of the fluorescent bacteria ratio. Calculation involved dividing each control/treated sample by the average value of the respective control/treated empty vector (EV). For example, the value for each biological replicate of the 184 kb reporter strain control was divided by the average value of biological samples from the EV control. Similarly, the value for each biological replicate of the 184 kb reporter strain supplemented with algal filtrates was divided by the average value of biological samples from the EV control supplemented with algal filtrates.

### Genetic manipulation of Roseobacter bacteria

To insert an antibiotic resistance gene into the 65 kb plasmid of *P. inhibens* DSM 17395, the mutant strain ES172 (Table S1) was generated by inserting a kanamycin-resistant gene in an intergenic region between gene PGA1_RS20320 and gene PGA1_RS19930. Regions of approximately 1000 bp upstream and downstream of the intergenic region were PCR-amplified using primers 1169, 1170, 1171 and 1172, respectively (Table S4). The amplified fragments were assembled and cloned into the pYDR8 vector (Table S6) using restriction-free cloning^43,44^. The resulting plasmid (pYDR9, Table S6) was introduced into competent *P. inhibens* by electroporation. Preparations of competent cells was performed as previously described^45^, with slight modifications. Shortly, cells were grown to an OD_600_ of 0.7 in ½ YTS with 40 g/L of sea salts (also termed full salts medium). Bacteria were washed 3 times using 10% (v/v) ice-cold glycerol, centrifuged each time for 5 min at 4 °C and 5,500 g. The competent cells were subsequently adjusted to an OD_600_ of 20 using 10% (v/v) ice-cold glycerol, frozen in liquid nitrogen and stored at -80°C. Electroporation was conducted with 300 μl aliquot of electrocompetent cells, using 10 µg of DNA in a 2 mm cuvette at 2.5 V, followed by 4 h recovery in ½ YTS with full salts. The transformed cells were then plated on ½ YTSS plates containing 150 μg/ml kanamycin, and resistant colonies were validated by PCR and DNA sequencing.

### Attachment assay

Bacterial cultures were diluted to an optical density of 0.01 at 600 nm (OD_600_) using 150 µl of 75% fresh algal filtrates diluted in 25% of ASW (v/v) for treatment and ASW for control. Both control and treated cultures were supplemented with L1 trace elements, L1 nutrients, vitamin mix, 33 mM Na_2_SO_4_, and 5 mM NH_4_Cl and 5.5 mM Glucose. The final control and treatment medium is referred to as CNS, with the treatment containing 75% volume of algal filtrates and 25% volume of ASW (v/v), in contrast to the control which consists solely of ASW. The diluted cultures were dispensed into 96-well flat-bottom plates to initiate the assay. To minimize evaporation during incubation, the plates were securely covered with UV-treated Parafilm. The bacterial cultures were incubated at 30 °C for 48 hours. The attachment assay was carried out following established protocols^46,47^. After incubation, the plates were subjected to the following steps: The culture medium was carefully aspirated, and the wells were rinsed twice with 1× phosphate-buffered saline (PBS, obtained from Sartorius, Beit HaEmek, Israel). This process aimed to remove non-adherent bacterial cells. The plates were dried by exposing them to a temperature of 55 °C for 20 minutes. The dried plates were submerged in a solution of 0.1% crystal violet. The plates were then incubated at room temperature for 10 minutes to allow the crystal violet to interact with the attached cells. The staining solution was discarded, and the plates were washed twice with PBS to remove excess crystal violet. The plates were left to dry overnight at room temperature to ensure complete fixation of the stained cells. To quantify the attached cells, the remaining crystal violet was extracted by adding 200 µL of 33% acetic acid to each well. The plates were incubated at room temperature for 15 minutes to facilitate extraction. Absorbance measurements were conducted at a wavelength of 595 nm using the TECAN microplate reader. The absorbance values were normalized to a blank consisting of the same growth medium, without bacterial cells, that was subjected to the same protocol.

### Generating the *Roseobacteraceae* T4SS dataset

All *Roseobacteraceae* genomes (of any quality) available in the Integrated Microbial Genomes & Microbiomes (IMG/M) database^48^ were scanned for the presence of the *virB8* gene using the IMG “Function search” tool and K03202 KEGG (Kyoto Encyclopedia of Genes and Genomes^49^) Orthology (KO) designation for VirB8. The *virB8* genes (K03203) were found in 320 out of 898 *Roseobacteraceae* genomes. The respective scaffolds were scanned for the presence of other *vir* genes (*virB1-B7, virB9-B11* and *virD4*) and were assessed as representing either plasmids or chromosomes (Table S7). Within these 320 genomes a total of 564 *virB8* genes were located on 553 scaffolds. Further assignment of the scaffolds to plasmids or chromosomes was conducted using the presence or absence of plasmid operational taxonomic unit (pOTU) designation that was assigned during genome annotation in IMG and according to scaffold size. Scaffolds with no operational taxonomic unit designation larger than 300 kb were marked as chromosomes. Scaffolds with no operational taxonomic unit designation and smaller than 300 kb were marked as unknowns. Of note, the T4SS is an extremely versatile apparatus. Conjugative T4SSs typically consist of the archetypal VirB/VirD4 components (encoded by *virB1*-*11* together with *virD4*). Yet, the precise number of needed genes and their essentiality for a functional T4SS remain poorly understood, particularly in non-model bacteria^50–53^. Consequently, due to the difficulty in predicting which T4SS is conjugative, we set stringent criteria and defined a potentially functional conjugative T4SS (*i.e.,* complete T4SS) as one that includes all the *vir* genes that comprise the T4SSs located on the conjugative plasmids of *D. shibae* and *P. inhibens* P72. Accordingly, T4SSs homologous to Vir but possessing other KO assignments (e.g., Tra), that were not experimentally shown to be conjugative in Roseobacters, were not included in this analysis. In total, our analysis identified 387 out of the 553 scaffolds as carrying a complete set of T4SS genes. However, it is possible that partially complete T4SSs in our analysis or homologous T4SSs (e.g., Tra) may be functional as well^54^.

## Results

### Establishing a conjugation assay using different Roseobacters

To monitor conjugation among various Roseobacters, we examined the ability of four different bacterial species - *Phaeobacter inhibens* P72, *Dinoroseobacter shibae, Marinovum algicola*, and *Phaeobacter inhibens* DSM 17395 - to act as donors and recipients in conjugation assays. These strains have been widely studied by us^25,55^ and by others^22,56–61^ as representative Roseobacters that specializes in interactions with microalgae. To establish a conjugation assay that can be followed in the lab, each strain was labeled with an antibiotic resistance gene on either its native plasmid or chromosome, allowing us to track the different participants using selection on antibiotics (Fig. 1A).

First, the *P. inhibens* P72 strain, carrying two complete T4SSs on both its 184 kb and 57 kb native plasmids (Fig. S1), was assessed as a donor. For the conjugation assays, the 57 kb plasmid was marked with a kanamycin-resistance gene and mated with three recipient strains: *P. inhibens* DSM 17395, *D. shibae*, and *M. algicola*, which were marked with a gentamicin-resistance gene on their chromosome (Fig. 1B). Second, the *D. shibae* strain, carrying two complete T4SSs on both its 126 kb and 191 kb native plasmids (Fig. S1), was evaluated as a donor. For the conjugation assays, the 126 kb plasmid was marked with a gentamicin-resistance gene and mated with two recipient strains: *P. inhibens* P72 and *P. inhibens* DSM 17395, both marked with a kanamycin-resistance gene on their chromosome (Fig. 1C). Following mating and selection, all assays resulted in colonies that exhibit resistance towards both kanamycin and gentamicin. To confirm that the double-resistance colonies were indeed transconjugants, several colonies were randomly chosen from each donor-recipient pair and the presence of the acquired plasmid, and the recipient chromosome was validated via PCR reactions. All tested transconjugants were confirmed to be the recipient strains that acquired the 57 kb plasmid (Fig. S2). Of note, several of these donor-recipient pairs were previously reported to execute successful conjugation under different experimental conditions^22,23^. Finally, we evaluated the ability of *P. inhibens* DSM 17395 to act as a donor. The *P. inhibens* DSM 17395 strain harbors three native plasmids of 65 kb, 78 kb, and 262 kb, and carries a complete T4SS on its chromosome (Fig. S1). We used three strains in which each of the native plasmids is marked with a kanamycin-resistance gene. The three strains were used as donors in conjugation assays with either *D. shibae* and *M. algicola* as recipients that were marked with a chromosomal gentamicin-resistance gene (Fig. 1D). Our results show that under our experimental conditions *P. inhibens* DSM 17395 is not capable of transferring any of its plasmids to recipient bacteria. To further support this result, we repeated the conjugation assays on agar plates instead of liquid media (Fig. S3), as previous studies demonstrated enhanced conjugation efficiency on solid surfaces like agar plates^13,62^. However, *P. inhibens* DSM 17395 was not capable of transferring any of its plasmids to recipient bacteria during conjugation on agar plates. As control, all experiments were accompanied by conjugation assays with a donor-recipient pair that was previously shown to result in successful conjugation^22^. Our data thus demonstrated that the antibiotic resistance-marked plasmids of *P. inhibens* P72 and *D. shibae* can be successfully transferred via conjugation to various Roseobacter recipients.

**Fig. 1.**
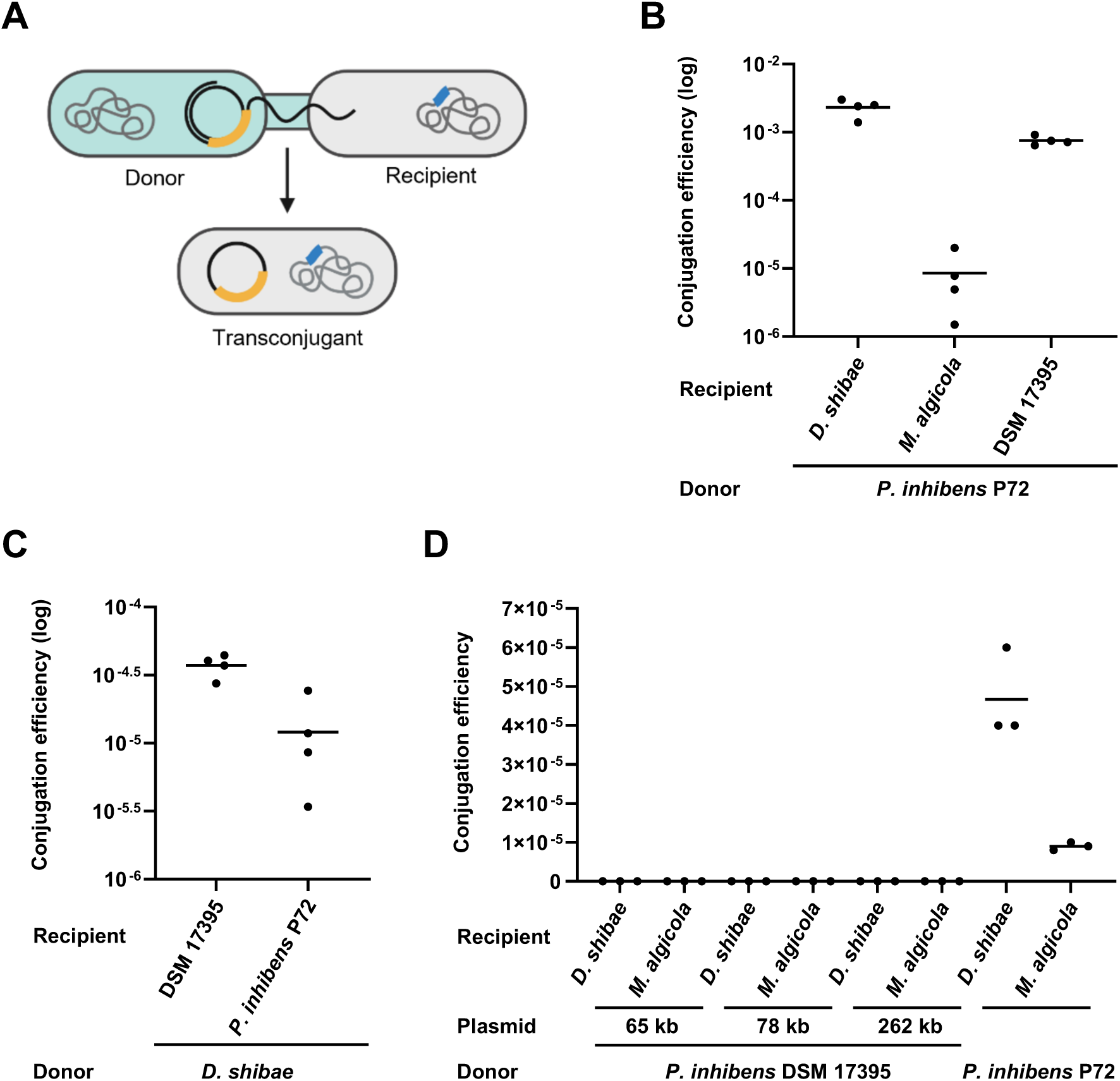
Establishing a robust conjugation assay. A) Schematic representation of the conjugation assay. The donor strain (light blue) carries an antibiotic resistance gene (yellow) on a native plasmid, while the recipient strain (grey) harbors a chromosomal antibiotic resistance gene (blue). Successful plasmid transfer from the donor to the recipient will result in transconjugants that are resistant to both antibiotics. **B-D)** Results of conjugation assays in liquid medium performed using different donor strains. **B)** *P. inhibens* P72 ^p57-kana^ served as a donor strain, carrying a kanamycin-resistance gene on its native 57 kb plasmid. The recipient strains *M*. *algicola* ^chr-gent^*, D. shibae* ^chr-gent^ and *P. inhibens* DSM 17395^chr-gent^, were marked with a gentamicin-resistance gene on their chromosome. **C)** *D. shibae* ^p126-gent^ served as a donor strain, carrying a gentamicin-resistance gene on its native 126 kb plasmid. Recipient strains were *P. inhibens* DSM 17395^chr-kana^ and *P. inhibens* P72 ^chr-kana^, both marked with a kanamycin-resistance gene on their chromosome. **D)** *P. inhibens* DSM 17395 served as a donor strain, carrying a kanamycin-resistance gene on its native 65 kb, 78 kb or 262 kb plasmids (*P. inhibens* DSM 17395 ^p65-gent^, *P. inhibens* DSM 17395 ^p78-gent^ and *P. inhibens* DSM 17395 ^p262-gent^ respectively). Recipient strains were *M*. *algicola* ^chr-gent^ and *D. shibae* ^chr-gent^, marked with a gentamicin-resistance gene on their chromosome. As a control, *P. inhibens* P72 ^p57-kana^ carrying a kanamycin-resistance gene on its native 57 kb plasmid was used as a donor and mated with the same recipients. Dots represent individual biological replicates, and lines indicate the mean values. Conjugation efficiency was calculated by normalizing transconjugant cell numbers to donor cell numbers. DSM 17395 indicates the *P. inhibens* DSM 17395 strain.

### Algal exudates enhance bacterial conjugation

Next, we investigated whether bacterial conjugation is influenced by proximity to an algal cell. To distinguish between proximity and the possible impact of algae as a surface for attachment, we chose to expose bacteria to algal exudates and not to actual algae. This approach allowed us to mimic the “chemical presence” of algae and probe its influence on conjugation between Roseobacter bacteria. We performed conjugation assays in which bacteria were treated with algal filtrates and compared them to control assays with untreated bacteria. Both treated and control assays were supplemented with essential nutrients to support bacterial growth. The conjugation assays were performed with *P. inhibens* P72 and *D. shibae* as plasmid donors to the recipients *D. shibae, M. algicola*, and *P. inhibens* DSM 17395 (Fig. 2). Our results show that conjugation assays that were supplemented with algal filtrates resulted in increased conjugation efficiency compared to untreated bacteria.

### Algal exudates enhance conjugation efficiency beyond the bacterial growth-promoting effect

Algal exudates contain metabolites that are known to promote bacterial growth^24,31,63^. To understand whether enhanced conjugation is merely the result of increased bacterial growth, we monitored the growth of donor strains, recipient strains, and transconjugants in pure cultures that were supplemented with algal filtrates, and compared them to untreated control cultures (Fig. 3A). Both treated and control cultures were supplemented with essential nutrients to support bacterial growth. Our data show that most bacteria displayed a shorter lag phase upon treatment with algal filtrates, a phenomenon previously documented by our laboratory^64^. Furthermore, although the assays were performed in nutrient replete conditions, several strains exhibited higher yields upon treatment with algal exudates.

Now, we aimed to determine whether algal filtrates enhance conjugation efficiency beyond their impact on bacterial growth dynamics. We therefore calculated the impact of algal filtrates on the growth of donors and recipients and compared it to the impact on growth of the transconjugants (Table 1). Our analysis demonstrate that algal exudates enhanced transconjugants cell numbers beyond the exudate impact on both donors and recipients, across all mated pairs. These findings suggest that algal filtrates enhance conjugative plasmid transfer beyond the growth promoting effect.

**Fig. 2.**
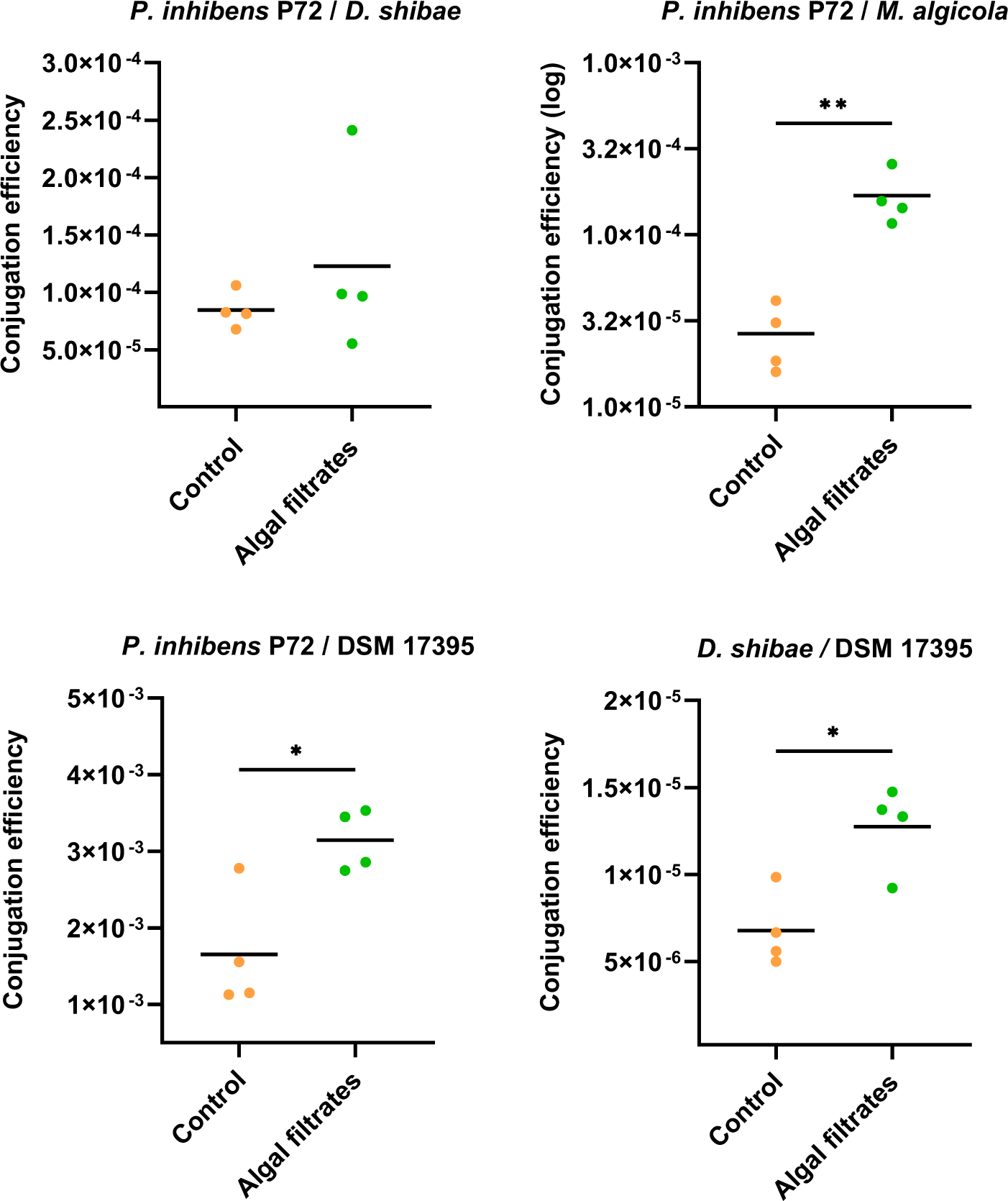
Algal exudates increase conjugation efficiency. Conjugation assays in liquid medium were performed using either *P. inhibens* P72 ^p57-kana^ as a donor, marked with a kanamycin-resistance gene on its native 57 kb plasmid, or *D. shibae* ^p126-gent^ as a donor, marked with a gentamicin-resistance gene on its native 126 kb plasmid. For *P. inhibens* P72 ^p57-kana^, three strains were tested as recipients: *M*. *algicola* ^chr-gent^*, D. shibae* ^chr-gent^ and *P. inhibens* DSM 17395^chr-gent^, all marked with a gentamicin-resistance gene on their chromosome. For *D. shibae* ^p126-gent^, the recipient strain was *P. inhibens* DSM 17395^chr-kana^, chromosomally marked with a kanamycin-resistance gene. Mating was performed with cultures treated under control conditions in CNS medium (orange dots) or with algal filtrates (green dots). Conjugation efficiency was calculated by normalizing transconjugant cell numbers to donor cell numbers. The title above each graph denotes *Donor/Recipient*. Dots represents individual biological samples, and lines indicate the mean values. Statistical significance was calculated using a t-test, with p<0.05 marked as *, and p<0.01 marked as **. DSM 17395 indicates the *P. inhibens* DSM 17395 strain.

**Fig. 3.**
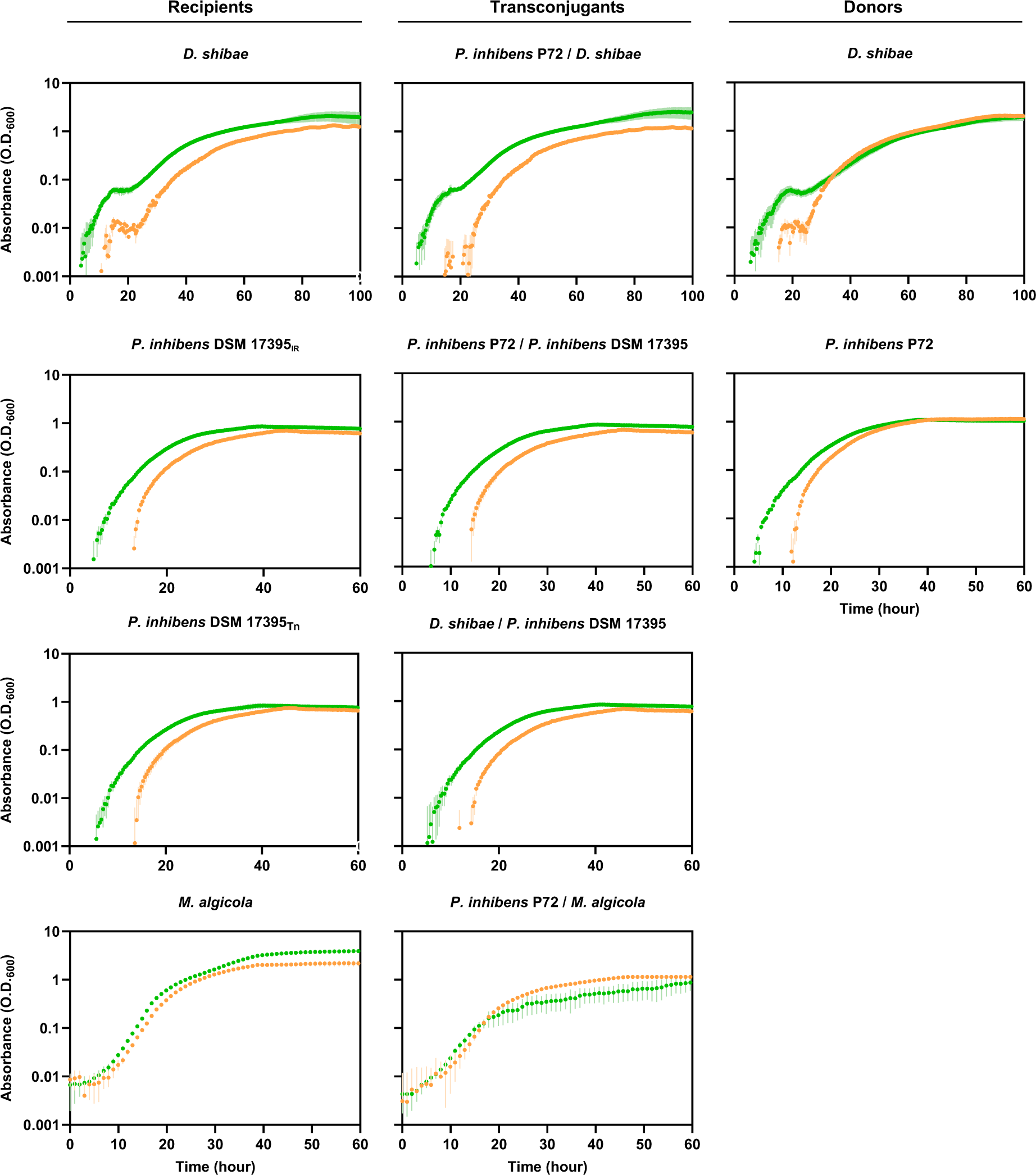
Algal filtrates effect on bacterial growth. Bacterial cultures of recipients (left column), transconjugants (middle column) and donors (right column), were treated with either CNS medium as control conditions (orange dots) or with algal filtrates (green dots). Growth was evaluated by measuring the optical density using a wavelength of 600 nm (O.D.600). Each dot represents the average of three biological replicates. The title above the column of transconjugants denotes *Donor/Recipient*.

**Table 1.**
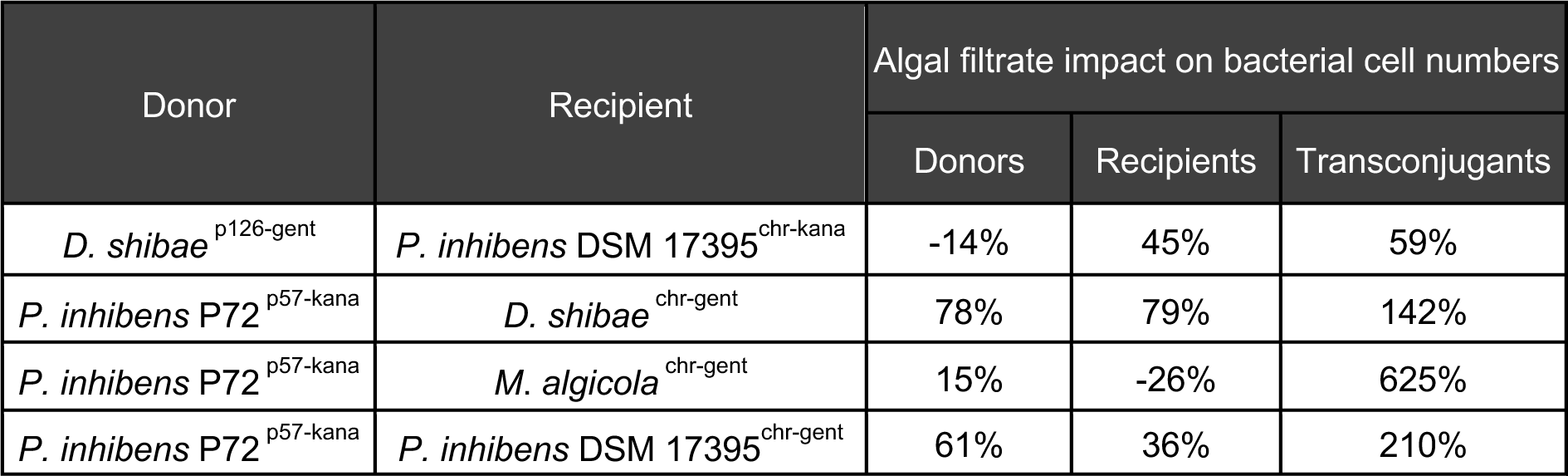
Algal filtrates enhance conjugation efficiency beyond bacterial growth promotion. The effect of algal filtrates on the growth of donors and recipients was calculated and compared to that of the transconjugants. Percentages were calculated by averaging colony forming units (CFU) numbers of donors, recipients and transconjugants separately (Table S8), for each conjugation pair (Fig. 2). Then, by dividing the averaged CFU numbers under algal filtrate treatment to the control, we determined the increase induced by the treatment in each group. For example, *P. inhibens* P72 ^p57-kana^ demonstrated 78% increase in CFU numbers due to the algal filtrates. As can be seen, the increase in in transconjugants numbers exceeded the increase observed for both donors and recipients across all conjugation pairs.

### Expression of the conjugation machinery is not influenced by algal exudates

To understand the mechanism underlying enhanced bacterial conjugation in response to algal exudates, we monitored the expression of the bacterial T4SS genes. The T4SS is mainly encoded by the *virB* operon while the *virD* operon carries several additional genes necessary for proper functioning of the T4SS^11,65^. To evaluate the impact of algal exudates on the expression of the T4SS, we analyzed expression levels in the donor strains *P. inhibens* P72 and *D. shibae* using qRT-PCR with primers that target the T4SS central genes *virB4*, *virB8* and *virD4.* We compared expression levels of bacterial cultures that were supplemented with algal filtrates versus non-treated cultures. Both treated and control cultures were supplemented with essential nutrients to support bacterial growth.

The results demonstrated that gene expression levels are similar between untreated cultures and cultures that were supplemented with algal filtrates, for both bacterial strains. Only expression of the *virD4* gene in *P. inhibens* P72 treated with algal filtrates showed slightly higher expression levels than untreated cultures (Fig. 4A). Additionally, the expression levels of *virB4*, *virB8* and *virD4* were comparable to those of the housekeeping genes, suggesting that under experimental conditions that support conjugation, the T4SS genes are expressed in a basal level but are not upregulated (Fig. S4).

Due to the rarity of conjugation events^66,67^, we questioned whether a qRT-PCR analysis, which averages RNA from the entire population, would possess the necessary sensitivity to detect such infrequent incidents. Instead, we aimed to observe the process at a single-cell resolution. Therefore, we developed reporter strains that contain a *gfp* gene under the *virB* promoter (Fig. 4B). However, analysis of these reporter strains using flow cytometry, following exposure to algal exudates, did not reveal increased expression (Fig. 4C). Taken together, these results suggest that algal-secreted compounds do not affect the expression of the T4SS machinery.

**Fig. 4.**
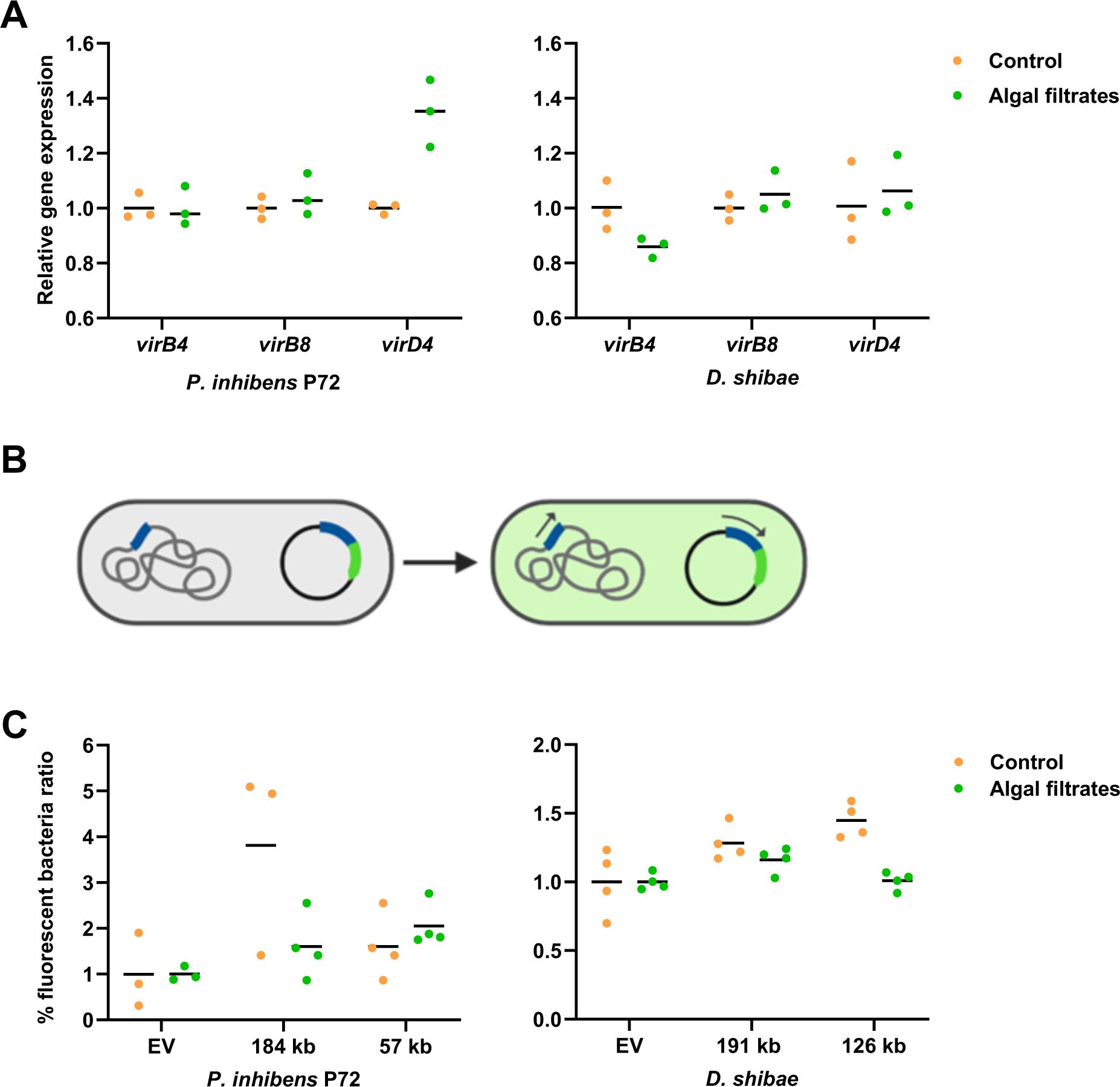
Algal exudates do not influence the expression of genes encoding the T4SS. A) Relative gene expression of *virB4*, *virB8* and *virD4* genes was assessed using qRT-PCR. RNA was extracted from bacterial cultures of *D. shibae* ^p126-gent^ and *P. inhibens* P72 ^p57-kana^ donor strains following treatment with either CNS medium as control conditions (orange dots) or with algal filtrates (green dots). RNA was extracted when bacterial cultures reached OD600 of 0.3. Each data point represents a biological replicate, lines represent the mean values of biological replicates. **B)** Schematic representation of the T4SS transcriptional reporter strains. The promoter of the *virB* operon (blue) was fused to GFP (green), cloned into a replicative plasmid, and transformed into *D. shibae* and *P. inhibens* P72. Activation of the native promoter is expected to lead to activation of the reporter construct, resulting in the expression of GFP and emission of a fluorescent signal. **C)** Reporter strains were generated using the T4SS promoters of the 57 kb and 184 kb native plasmids of *P. inhibens* P72, and the 126 kb and 191 kb native plasmids of *D. shibae.* Bacteria carrying these constructs were supplemented either with CNS as a control (orange dots) or algal exudates (green dots). Dots represent the percentage of fluorescent bacteria. For each biological sample, the value was normalized to the average of its corresponding control (see Methods section). Each dot represents a biological replicate, and lines represent the mean values.

### Algal exudates increase bacterial attachment capabilities

In the absence of increased expression of T4SS genes upon exposure to algal exudates, the mechanism behind the enhanced bacterial conjugation remains unclear. We hypothesized that enhanced conjugation might be facilitated by increased bacterial adhesion. Previous research in our laboratory has shown that algal exudates enhance the attachment of wild-type bacteria from the *P. inhibens* DSM 17395 strain^46^. Now, we sought to investigate whether similar effects are observed when the diverse Roseobacters in our current study are exposed to exudates (Fig. 5). Using an attachment assay, we evaluated the adhesion capabilities of the various donors and recipients, and tested whether algal exudates affect their adhesion. Our results show that algal exudates significantly improve the attachment capabilities of *P. inhibens* P72 and *P. inhibens* DSM 17395 strains. *D. shibae* and *M. algicola* in contrast, do not appear to have attachment capabilities in our experimental conditions. Of note, in all our conjugation assays, either the donor or recipient strain possessed attachment capabilities that are enhanced following exposure to algal exudates. Therefore, increased attachment could explain the enhanced conjugation efficiency.

**Fig. 5.**
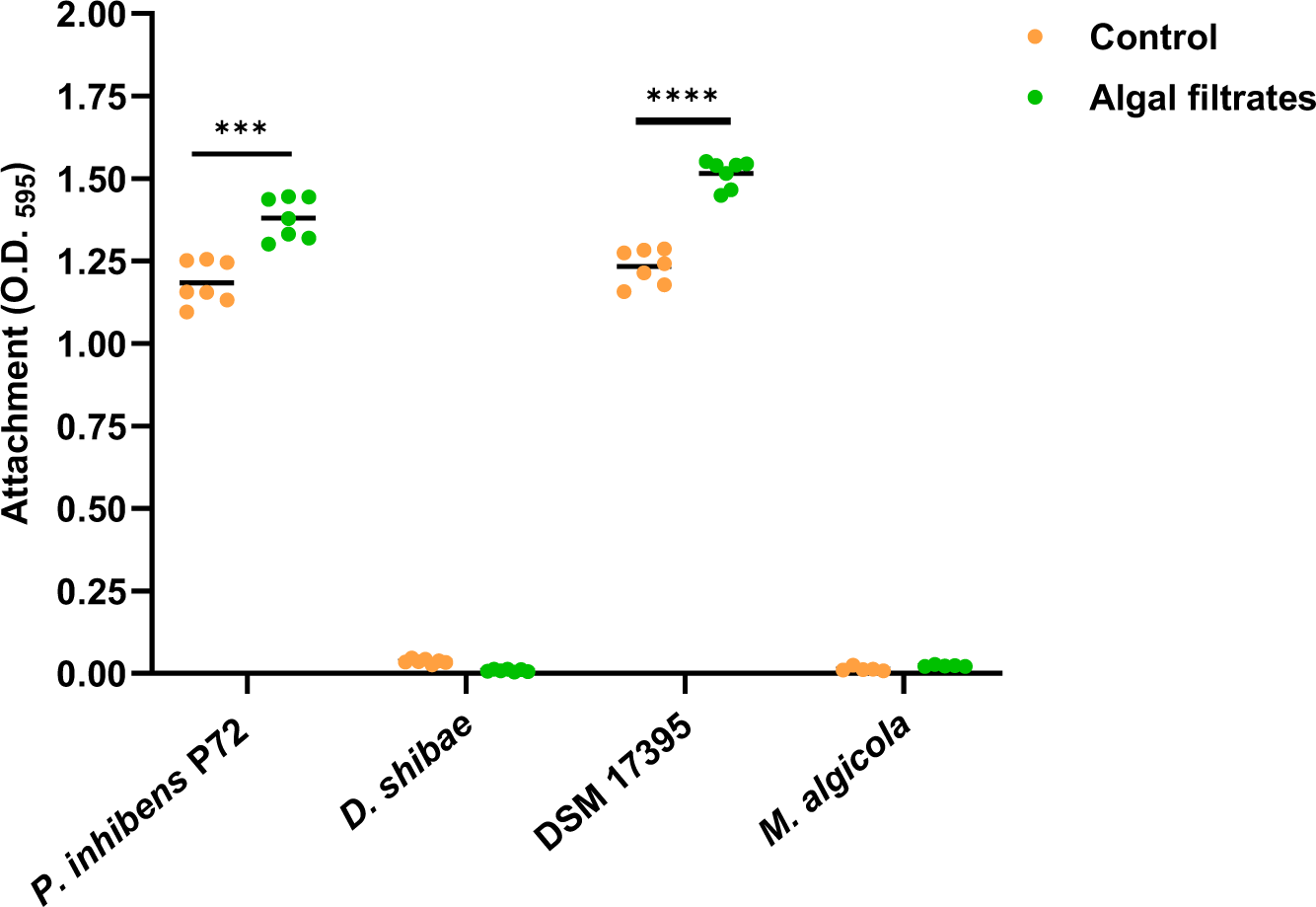
Algal exudates promote bacterial surface attachment. Bacterial donor and recipient strains were treated either with CNS as control (orange dots) or with algal filtrates (green dots). The bacterial donor strains were *P. inhibens* P72 ^p57-kana^ marked with a kanamycin-resistance gene on its native 57 kb plasmid and *D. shibae* ^p126-gent^ marked with a gentamicin-resistance gene on its native 126 kb plasmid. The recipient strains were *P. inhibens* DSM 17395^chr-gent^ and *M*. *algicola* ^chr-gent^ marked with a gentamicin-resistance gene on the chromosome. Attachment was assessed by measuring the absorbance at O.D.595 of the Crystal Violet dye from bacterial cells that were strongly attached to 96-well plates following an attachment assay (see Methods section). Each sample was normalized to Crystal Violet extracted from the same medium but without bacteria. Each strain and treatment consisted of n=7 wells. Dots represent individual biological replicates, and lines indicate the mean values. Statistical significance was calculated using a t-test, with 0.0001<p<0.001 marked as ***, and p<0.0001 marked as ****.

### Conjugation enhancement by algal exudates is reduced when conducted on solid surface

We further investigated the correlation between enhanced conjugation and heightened bacterial attachment triggered by algal exudates. To understand if the enhanced conjugation efficiency resulted from increased attachment, we conducted the conjugation assay under conditions where attachment no longer conferred an advantage. Thus, we performed conjugation assays on agar plates using a dense, thoroughly mixed inoculum of both donors and recipients. Unlike in liquid-based conjugation assays where bacteria swim and locate their conjugation partners, assays on plates facilitate immediate contact between all bacteria, thereby eliminating the advantage of attachment^13,62^. The conjugation assays on plates were conducted using *P. inhibens* P72 and *D. shibae* as plasmid donors, while recipients included *D. shibae, M. algicola*, and *P. inhibens* DSM 17395 (Fig. 6). Assays were treated with algal filtrates and compared to untreated conditions, and all assays were supplemented with essential nutrients to support bacterial growth. Our findings reveal that the majority of conjugation assays supplemented with algal filtrates exhibited conjugation efficiencies comparable to the control. Thus, when attachment no longer confers an advantage, the impact of algal exudates on bacterial conjugation is reduced. These results support the role of algal exudates in promoting bacterial attachment, thus facilitating enhanced conjugation.

**Fig. 6.**
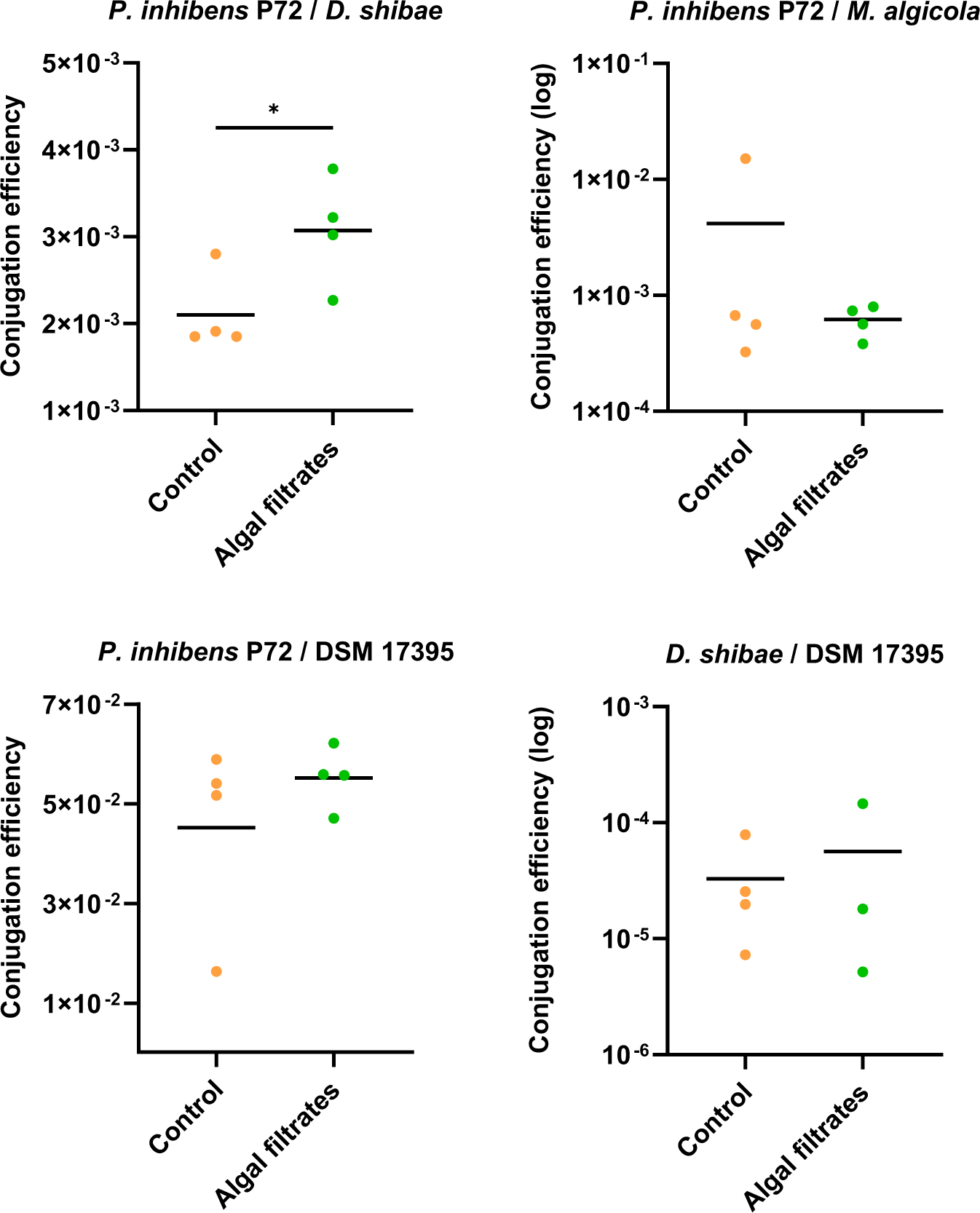
Algal exudates exhibit a lower impact on conjugation when performed on agar plates. Conjugation assays on agar plates were performed using either *P. inhibens* P72 ^p57-kana^ as a donor, marked with a kanamycin-resistance gene on its native 57 kb plasmid, or *D. shibae* ^p126-gent^ as a donor, marked with a gentamicin-resistance gene on its native 126 kb plasmid. For *P. inhibens* P72 ^p57-kana^, three strains were tested as recipients: *M*. *algicola* ^chr-gent^*, D. shibae* ^chr-gent^ and *P. inhibens* DSM 17395^chr-gent^, all marked with a gentamicin-resistance gene on their chromosome. For *D. shibae* ^p126-gent^, the recipient strain was *P. inhibens* DSM 17395^chr-kana^, chromosomally marked with a kanamycin-resistance gene. Mating was performed with cultures treated under control conditions with CNS medium (orange dots) or with algal filtrates (green dots). Conjugation efficiency was calculated by normalizing transconjugant cell numbers to donor cell numbers. The title above each graph denotes *Donor/Recipient*. Dots represents individual biological samples, and lines indicate the mean values. Statistical significance was calculated using a t-test, with p<0.05 marked as *. DSM 17395 indicates the *P. inhibens* DSM 17395 strain.

## Discussion

### HGT and the ecological success of Roseobacters

The current study reveals that algal exudates increase bacterial conjugation, and consequently HGT, among Roseobacter bacteria. HGT has been suggested to play important roles in the resilience of marine bacteria^68^ and is considered key in the ecological abundance and success of Roseobacters^69–71^. Various mechanisms for HGT have been observed in Roseobacters, including GTAs^68^ and conjugation^21^. Moreover, Roseobacters have many native plasmids^70,71^, up to a dozen plasmids in a single bacterium^70^. Many of these plasmids harbor T4SSs^13^ that may facilitate conjugative HGT, potentially supporting rapid adaptation. Indeed, 27% of sequenced Roseobacters genomes harbor all genes needed for a functional conjugative T4SS (Fig. S5). Conjugation initiation and the successful transfer of DNA was shown to be impacted by factors such as signaling molecules, temperature, cell physiology, population density, pH, and nutrients^13,66,72^. However, in marine bacteria, little is known about the conditions required to induce HGT, particularly conjugation. Here, we show that algal secreted metabolites promote HGT in Roseobacters.

### Bacterial HGT in the algal phycosphere

In marine ecosystems, algae provide bacteria with nutrients^24,31,63^ and an attachment surface^25,32^. By attaching to an algal cell, bacteria can secure a steady flux of algal exudates in the algal phycosphere, which is the immediate volume surrounding the algal cell. Indeed, Roseobacter bacteria often co-occur with various algal hosts in the environment and in laboratory cultures^25,38,73–75^. As multiple bacteria can attach onto an algal host, they may engage in processes that require cell-to-cell contact, such as conjugation. Therefore, proximity and attachment onto an algal host may increase the chances for plasmid transfer events among bacteria in the algal phycosphere^13,21,34,35^. Interestingly, environmental experiments with phytoplankton-derived dissolved organic matter demonstrated attraction of bacteria harboring T4SS genes, as shown by metagenomic analysis^76^. Moreover, algal exudates support bacterial growth and metabolism^77^, further enhancing bacterial density, which in turn could augment the chances of bacterial HGT.

### Algal exudates increase attachment and enhance HGT

HGT mechanisms rely on proximity, with conjugation specifically requiring cell-to-cell contact^78^. Therefore, conditions that stabilize and facilitate cell-to-cell contact will enhance the opportunities for conjugation events^20^. Indeed, conjugation in biofilms was extensively studied^13^. Conjugation is enhanced in biofilms, and further stimulates biofilm development^14–19^. Specifically, it was shown that factors mediating formation and stabilization of cell-to-cell contacts in biofilms promote T4SS mediated gene transfer^20^. Additionally, attachment to abiotic surfaces was also demonstrated to increased conjugation^79^. Recent studies have revealed that microalgae and Roseobacter bacteria create a shared algal-bacterial extracellular matrix, fostering algal-bacterial aggregation^46^. Algal exudates stimulate bacterial exopolysaccharide production, which contributes to the formation of a joint algal-bacterial extracellular matrix. Our data illustrate that algal exudates primarily enhance Roseobacter conjugation by promoting bacterial attachment. Consequently, it is plausible that algal-bacterial aggregates may facilitate enhanced conjugation, although further investigation is required to confirm this hypothesis.

### The T4SS genomic location

Establishing a conjugation assay resulted in characterizing Roseobacter strains capable of serving as donors in conjugation, as well as a strain with a chromosomally encoded T4SS unable to donate a plasmid during conjugation (Fig. 1D). Various bacteria carry the T4SS-encoding genes on plasmids, chromosomes, and even on both^80,81^. While plasmid-encoded T4SSs are known to commonly mediate conjugation^82^, chromosomally encoded T4SSs rarely mobilize a plasmid to a recipient strain^80^. Chromosomally encoded T4SSs are mainly involved in processes such as protein secretion, ssDNA secretion into the extracellular milieu, natural transformation, and symbiosis^54,58,81,83–89^. Of note, within sequenced Roseobacters genomes, 23% carry only plasmid-encoded T4SSs, 2% carry only chromosome-encoded T4SSs, and 1% carry T4SSs on both the chromosome and a plasmid (Fig. S5). Given the notable fraction of chromosome-encoded T4SSs in Roseobacters, the function of these secretion systems merits further exploration^58^.

### The algal impact on bacterial HGT and evolution

The evolution of marine heterotrophic bacteria is linked to their micro-algal hosts, as manifested by the many examples of algal-bacterial chemical exchanges that likely arose during their co-evolution^25–27,55,57,90^. The current study reveals the impact that algal exudates have on bacterial conjugation, potentially through bacterial attachment, which adds another layer to our understanding of bacterial evolution in the ocean. Algal exudates represent a distinct chemical signature of a given algal species^91^, and consequently, attract phylogenetically and functionally discrete populations of bacteria^76,77,92^. By attracting specific bacterial populations, algae have the potential to affect the bacterial partners that perform genetic exchange, thus possibly influencing bacterial evolution.

## Acknowledgments

We are grateful to Dr. Jӧrn Petersen (Leibniz Institute, DSMZ, Germany) for generously sharing many valuable bacterial strains. We thank Prof. Juan Barja (University of Santiago de Compostela, Spain) for kindly sharing the *P. inhibens* P72 bacterial strain. We thank all members of the Segev lab for insightful comments and discussions.

## Author contribution

YDR and ES designed the study. YDR, VL, LY, DM, and IVK performed and analyzed experiments. YDR and ES wrote the manuscript. All authors discussed the results and contributed to the final manuscript.

## Data availability

Strains and plasmids used in this study are listed in Tables S1 and S6, respectively. Engineered bacteria and plasmids are available from the corresponding author on reasonable request. Primers designed for bacterial identification and engineering and for qRT-PCR analysis, are available in Tables S4 and S5 respectively. Data used for supplementary Fig. S5 is presented in Table S7.

