## Supplemental Information for "Algal Exudates Promote Conjugation in Marine Roseobacters"

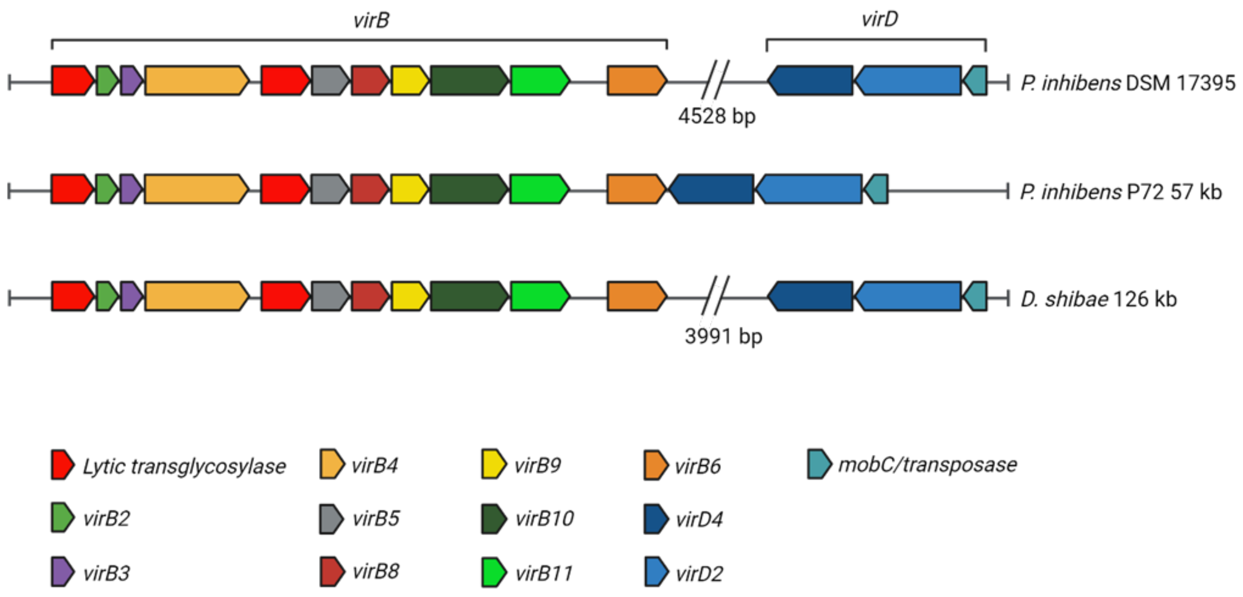


**Fig. S1. Genetic organization of the T4SS genes in the donor strains.** Schematic representation of genes from the *virB* and *virD* operons, which construct the T4SS, located in the chromosome of *P. inhibens* DSM 17395, the 57 kb plasmid of *P. inhibens* P72 and the 126 kb plasmid *of D. shibae*. Homologous genes are shown in the same color code. Of note, a potentially functional conjugative T4SS (*i.e.,* complete T4SS) was determined as one that includes all the *vir* genes that comprise the T4SSs located on the conjugative plasmids of *D. shibae* and *P. inhibens* P72.

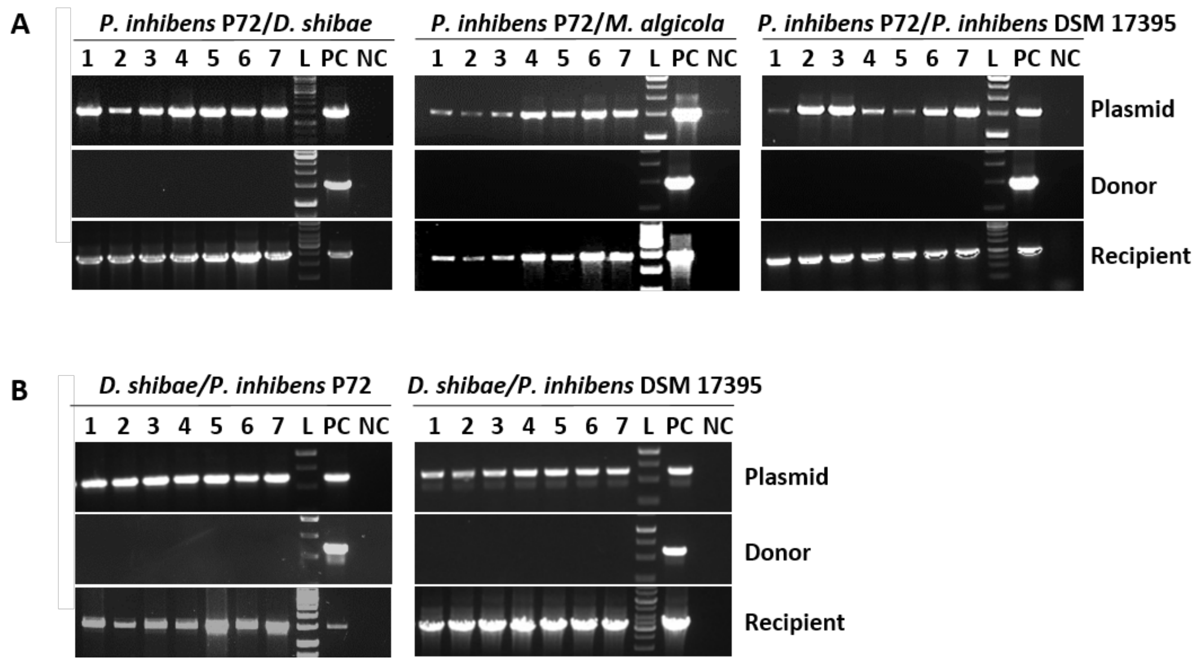


**Fig. S2. Validation of double-resistance colonies from conjugation assays as transconjugants.** Gel images of PCR products for validation of the transconjugants. **A)** *P. inhibens* P72$^{\text{p57-kana}}$ as a donor strain, *D. shibae* DFL-12$^{\text{chr-gent}}$, *M. algicola* DG898$^{\text{chr-gent}}$, and *P. inhibens* DSM 17395$^{\text{chr-gent}}$ as recipients**. B)** *D. shibae* DFL-12$^{\text{p126-gent}}$ as a donor strain, *P. inhibens* DSM 17395$^{\text{chr-kana}}$ as recipient. From each donor-recipient pair (Fig. 2), seven transconjugants colonies were randomly selected (marked 1-7). Each colony was streaked three times, followed by genomic extraction and PCR reactions. The transconjugants were tested using three sets of primers that identify the donor strain, recipient strain, and the transferred plasmid. Positive controls (PC) included primers and the corresponding strain DNA, and negative controls (NC) included the opposing strain DNA (i.e., primers targeting the donor strain with DNA of the recipient strain). The title above each graph denotes *Donor/Recipient*. L marks the 1 kb ladder incorporated in all gels

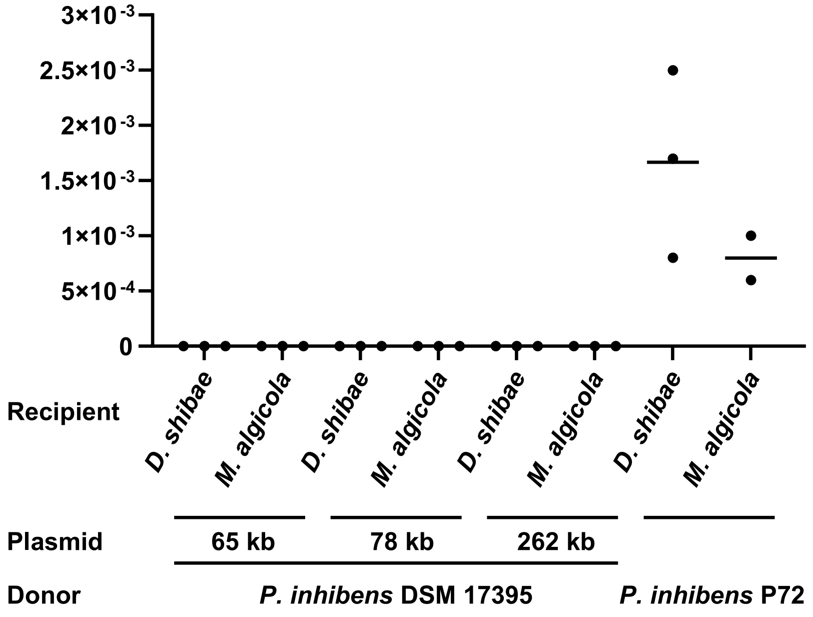


**Fig. S3. The *P. inhibens* strain DSM 17395 is unable to serve as a plasmid donor.** Results of conjugations assays performed on MB agar plates. Conjugation assays were carried out using *P. inhibens* DSM 17395 as the donor, with a kanamycin-resistance gene on one of its native 65 kb, 78 kb and 262 kb plasmids (*P. inhibens* DSM 17395 $^{p65\text{-}gent}$, *P. inhibens* DSM 17395 $^{p78\text{-}gent}$, and *P. inhibens* DSM 17395 $^{p262\text{-}gent}$ respectively). Each donor strain was mated with recipient strains *M. algicola* DG898 $^{chr\text{-}gent}$ or *D. shibae* DFL-12 $^{chr\text{-}gent}$, marked with a gentamicin-resistance gene on their chromosome. As a control, *P. inhibens* P72 $^{p57\text{-}kana}$, carrying a kanamycin-resistance gene on its 57 kb native plasmid was used as a donor and mated with the same recipients. Dots represent individual biological replicate and lines indicate the mean value. Conjugation efficiency was calculated by normalizing transconjugant cell numbers to donor cell numbers.

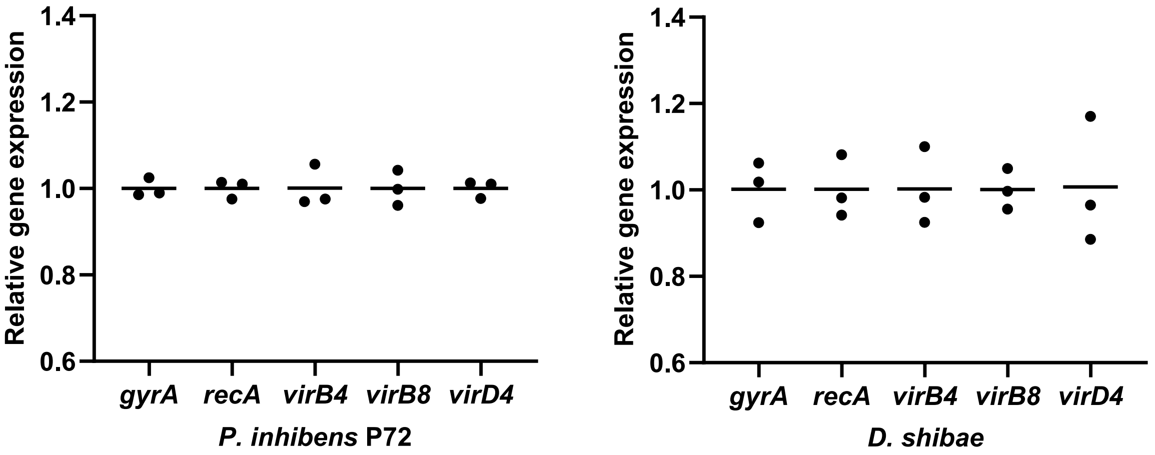


**Fig. S4. Expression of T4SS genes is comparable to those of the housekeeping genes.** Relative gene expression of *virB4*, *virB8* and *virD4* genes was assessed using qRT-PCR and compared to the house keeping genes *gyrA* and *recA*. RNA was extracted from bacterial cultures of *D. shibae* DFL-12 $^{chr\text{-}gent}$ and *P. inhibens* P72 $^{p57\text{-}kana}$ grown in CNS medium, when they reached $OD_{600}$ of 0.3. Data points represent the mean of three technical replicates originating from one biological sample, lines represent replicates mean values.

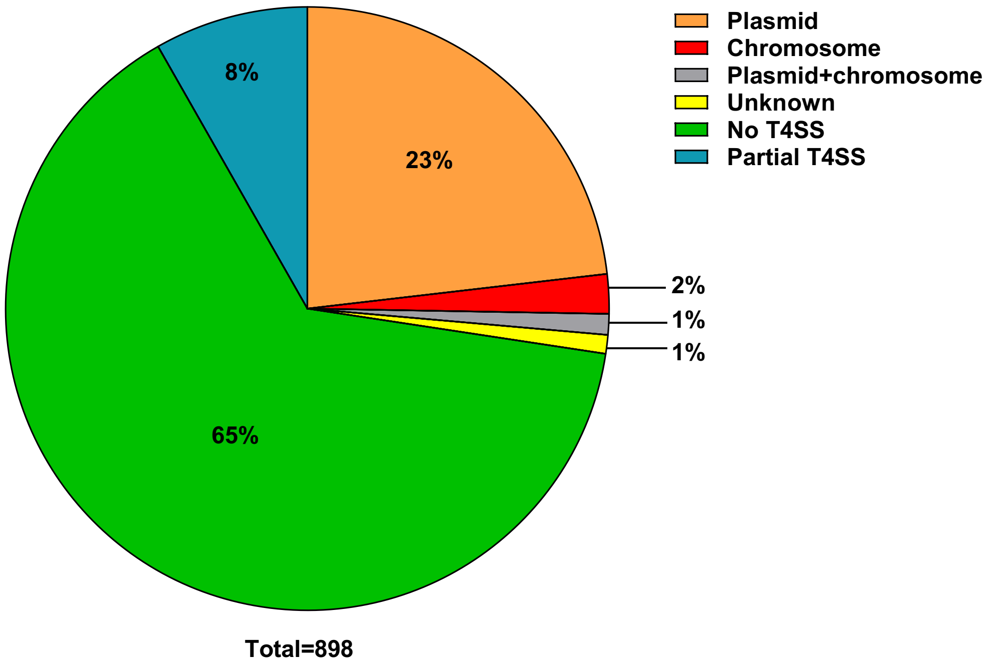


**Fig. S5. Chromosome-encoded and plasmids-encoded T4SSs in Roseobacter bacteria.** Roseobacter bacteria genomes were searched for T4SSs, and percentages were calculated for genomes with no T4SSs (green), partial T4SSs (blue), complete plasmid-encoded T4SSss (orange), complete chromosome-encoded T4SSs (red), complete plasmid-encoded and chromosome-encoded T4SSs located in the same genome (grey), and complete T4SSs with an unknown locus (yellow). Of note, a potentially functional conjugative T4SS (*i.e.,* complete T4SS) was determined as one that includes all the *vir* genes that comprise the T4SSs located on the conjugative plasmids of *D. shibae* and *P. inhibens* P72.

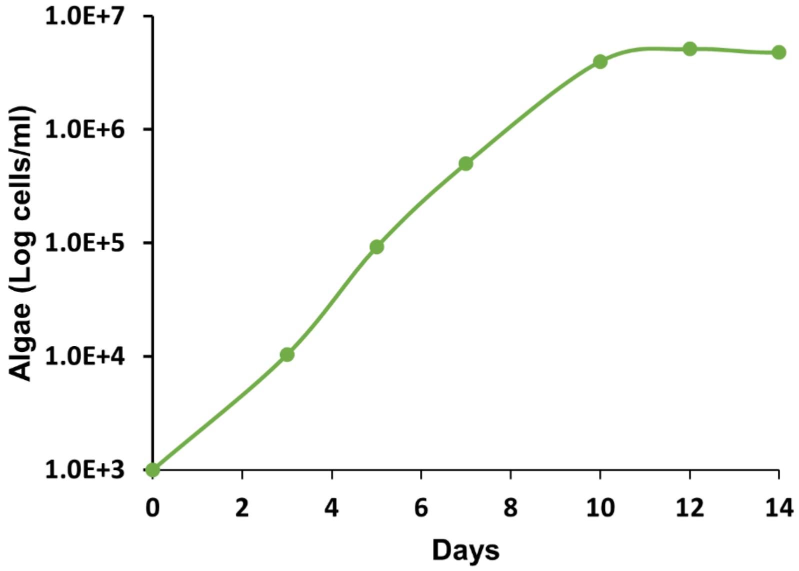


**Fig. S6. Growth curves of axenic *Emiliania huxleyi* cultures**. The cultures were cultivated at 18°C in ASW with L1 supplements, and monitored for 14 days, with spent media collected on day 12. Each data point in the growth curve represents the average from three biological replicates. Error bars indicate the standard deviation.

**Table S1:** List of bacterial strains and mutants.

| Strain | Tagged locus | Insertion site | Reference | Lab collection identifier | Resistance | Comments |
|---|---|---|---|---|---|---|
| *Phaeobacter inhibens* DSM 17395 | | | DSMZ, Germany | ES1 | | Wild type |
| *Phaeobacter inhibens* DSM 17395 | 65 kb plasmid | 18,622 | Current study | ES172 | Kanamycin | Intergenic region mutant. |
| *Phaeobacter inhibens* DSM 17395 | 78 kb plasmid | 26,249-30,284 | Lab collection | ES171 | Kanamycin | Knock-out mutant in both PGA1_RS19420 and PGA1_RS194 15. |
| *Phaeobacter inhibens* DSM 17395 | 262 kb plasmid | 138,002 | Dr. Jörn Petersen from the Leibniz Institute, DSMZ, Germany. | ES170 | Kanamycin | Insertion in PGA1_262p01260, generated with the <R6kyori/KAN-2> transposon-kit. |
| *Dinoroseobacter shibae* DFL-12 | | | DSMZ, Germany | ES9 | | Wild type |
| *Dinoroseobacter shibae* DFL-12 | 126 kb plasmid | 27,457 | Patzelt *et al.*[23] | ES94 | Gentamicin | Insertion in Dshi_3964, generated using mariner transposon mutagenesis. |
| *Dinoroseobacter shibae* DFL-12 | Chromosome | 3,502,458 | Birmes *et al.*[22] | ES95 | Gentamicin | Intergenic region mutant, generated using mariner transposon mutagenesis. |
| *Phaeobacter inhibens* P72 | | | Dr. Jörn Petersen from the Leibniz Institute, DSMZ, Germany and Prof. Juan Barja from University of Santiago de Compostela, Spain | ES98 | | Wild type |
| *Phaeobacter inhibens* P72 | 57 kb plasmid | 38,628 | Birmes *et al.*[22] | ES96 | Kanamycin | Insertion in CP010740, generated with the <R6kyori/KAN-2> transposon-kit. |
| *Phaeobacter inhibens* P72 | Chromosome | 1,678,662 | Dr. Jörn Petersen from the Leibniz Institute, DSMZ, Germany | ES168 | Kanamycin | Insertion in PhaeoP72_RS07995, generated with the <R6kyori/KAN-2> transposon-kit. |
| *Marinovum algicola* DG898 | Chromosome | 3,256,562 | Birmes *et al.*[22] | ES93 | Gentamicin | Insertion in MALG_03188, generated using |

| | | | | | | |
|---|---|---|---|---|---|---|
| | | | | | | mariner transposon mutagenesis. |
| *Phaeobacter inhibens* DSM 17395 | Chromosome | 1,400,112 | Lab collection | ES70 | Gentamicin | Intergenic region mutant. |
| *Phaeobacter inhibens* DSM 17395 | Chromosome | 2,372,826 | Dr. Jörn Petersen from the Leibniz Institute, DSMZ, Germany. | ES97 | Kanamycin | Insertion in PGA1_RS11385, generated with the <R6kyori/KAN-2> transposon-kit. |
| *Escherichia coli* ST18 DSM 22074 | | | DSMZ, Germany | ES201 | | |

**Table S2:** List of donor-recipient conjugation pairs. Donors are denoted by their transferred plasmid size in kilobases (Kb) represented as "p" for plasmid, and the antibiotic resistance gene that marks the plasmid ("gent" for gentamicin-resistance and "kana" for kanamycin-resistance). Recipients are designated as "chr" and the antibiotic resistance gene that marks their chromosome.

| Donor | Recipient |
|---|---|
| *P. inhibens* DSM 17395 $^{\text{p262-gent}}$ | *D. shibae* DFL-12 $^{\text{chr-gent}}$ |
| *P. inhibens* DSM 17395 $^{\text{p78-gent}}$ | *D. shibae* DFL-12 $^{\text{chr-gent}}$ |
| *P. inhibens* DSM 17395 $^{\text{p65-gent}}$ | *D. shibae* DFL-12 $^{\text{chr-gent}}$ |
| *P. inhibens* DSM 17395$^{\text{p262-gent}}$ | *M. algicola* DG898 $^{\text{chr-gent}}$ |
| *P. inhibens* DSM 17395 $^{\text{p78-gent}}$ | *M. algicola* DG898 $^{\text{chr-gent}}$ |
| *P. inhibens* DSM 17395 $^{\text{p65-gent}}$ | *M. algicola* DG898 $^{\text{chr-gent}}$ |
| *P. inhibens* P72 $^{\text{p57-kana}}$ | *D. shibae* DFL-12 $^{\text{chr-gent}}$ |
| *P. inhibens* P72 $^{\text{p57-kana}}$ | *M. algicola* DG898 $^{\text{chr-gent}}$ |
| *P. inhibens* P72 $^{\text{p57-kana}}$ | *P. inhibens* DSM 17395$^{\text{chr-gent}}$ |
| *D. shibae* DFL-12 $^{\text{p126-gent}}$ | *P. inhibens* DSM 17395$^{\text{chr-kana}}$ |
| *D. shibae* DFL-12 $^{\text{p126-gent}}$ | *P. inhibens* P72 $^{\text{chr-kana}}$ |

**Table S3:** Primers for transconjugant validation. Each transconjugant was validated using three primer sets identifying the donor, the recipient, and the transferred plasmid. Primer sequences are listed in table 5. Donors are denoted by their transferred plasmid size in kilobases (Kb) represented as “p” for plasmid, and the antibiotic resistance gene that marks the plasmid (“gent” for gentamicin-resistance and “kana” for kanamycin-resistance). Recipients are designated as “chr” and the antibiotic resistance gene that marks their chromosome.

| Strain | Target of the primer set | Primer name |
|---|---|---|
| *D. shibae* DFL-12 $^{p126\text{-}gent}$ | Donor | 802+803 |
| *P. inhibens* P72 $^{p57\text{-}kana}$ | Donor | 794+777 |
| *P. inhibens* DSM 17395$^{chr\text{-}kana}$ | Recipient | 126+92 |
| *D. shibae* DFL-12 $^{chr\text{-}gent}$ | Recipient | 802+803 |
| *M. algicola* DG898 $^{chr\text{-}gent}$ | Recipient | 804+805 |
| *P. inhibens* DSM 17395$^{chr\text{-}gent}$ | Recipient | 47+290 |
| *P. inhibens* P72 $^{chr\text{-}kana}$ | Recipient | 796+797 |
| *D. shibae* DFL-12, 126 kb plasmid | plasmid | 772+773 |
| *P. inhibens* P72, 57 kb plasmid | plasmid | 808+809 |

**Table S4:** Primer list for PCR reactions and construction of mutant strain. Chromosomally marked strains are designated as "chr". Antibiotic resistance markers are indicated by "gent" for gentamicin-resistance and "kana" for kanamycin-resistance.

| Primer name | Sequence | Strain | Comments | Origin |
|---|---|---|---|---|
| 47 | CAGGATGAGTTATTGCACCA | *P. inhibens* DSM 17395$^{chr\text{-}gent}$ | Validation of transconjugants | Current study |
| 290 | GAGCCTACATGTGCGAATGATGC | *P. inhibens* DSM 17395$^{chr\text{-}gent}$ | Validation of transconjugants | Current study |
| 92 | CCAGAGAACAATAGTAAGAGCG | *P. inhibens* DSM 17395$^{chr\text{-}kana}$ | Validation of transconjugants | Current study |
| 126 | GCTTGATGTCGAGGTGCTGT | *P. inhibens* DSM 17395$^{chr\text{-}kana}$ | Validation of transconjugants | Current study |
| 772 | GGCACCATCGTCGGAACCAAT | *D. shibae* DFL-12, 126 kb plasmid | Validation of transconjugants | Patzelt *et al.*[23] |
| 773 | TGGTATCAGGCATTCGCTTCA | *D. shibae* DFL-12, 126 kb plasmid | Validation of transconjugants | Patzelt *et al.*[23] |
| 777 | AGATCGCCAAATTGCAGCAG | *P. inhibens* P72 | Validation of transconjugants | Current study |
| 794 | AGACGACCCCTTGAGGCAC | *P. inhibens* P72 | Validation of transconjugants | Current study |
| 802 | ACGGCATCGACCACTTCTTC | *D. shibae* DFL-12 | Validation of transconjugants | Birmes *et al.*[22] |
| 803 | ACGGAAGATCGGGTACATGG | *D. shibae* DFL-12 | Validation of transconjugants | Birmes *et al.*[22] |
| 804 | CAATCCGATCGACAGCTACG | *M. algicola* DG898 | Validation of transconjugants | Birmes *et al.*[22] |
| 805 | TGATCGGCATGTAGAGCAGC | *M. algicola* DG898 | Validation of transconjugants | Birmes *et al.*[22] |
| 808 | GAACCAGGAATGCATCGGTC | *P. inhibens* P72, 57b kb plasmid | Validation of transconjugants | Birmes *et al.*[22] |
| 809 | CCTGATCCGGCTATGACCAT | *P. inhibens* P72, 57b kb plasmid | Validation of transconjugants | Birmes *et al.*[22] |
| 974 | GCGAATTTTAACAAAATATTAACG CTTACATATGTGCAGCCTTCTCAA CCTGAGG | *D. shibae* DFL-12 | 191 kb plasmid reporter strain | Current study |
| 975 | CACCAGTGAACAGCTCTTCGCCT TTACGCATCTGCGCATTTGCGGA ACCCGTCAGGAACCAG | *D. shibae* DFL-12 | 191 kb plasmid reporter strain | Current study |
| 972 | CGCGAATTTTAACAAAATATTAAC GCTTACACCAAACTTGCGCAGCG AGGAAGCGTCTC | *D. shibae* DFL-12 | 126 kb plasmid reporter strain | Current study |
| 973 | CACCAGTGAACAGCTCTTCGCCT TTACGCATTTGCGCAGTGGCGGG ACCCGCGAGATACCAG | *D. shibae* DFL-12 | 126 kb plasmid reporter strain | Current study |
| 987 | GCGAATTTTAACAAAATATTAACG CTTACATGCCACCCATCGGCTCA ACCTTAAGTTTCTG | *P. inhibens* P72 | 184 kb plasmid reporter strain | Current study |
| 988 | CACCAGTGAACAGCTCTTCGCCT TTACGCATCTGCGCATTGGCGGA ACCCGTCAGGAACCAG | *P. inhibens* P72 | 184 kb plasmid reporter strain | Current study |
| 989 | GCGAATTTTAACAAAATATTAACG CTTACAGTAGCCCGCTCTAGCCC GTTGGGTCATCCTG | *P. inhibens* P72 | 57 kb plasmid reporter strain | Current study |
| 990 | CACCAGTGAACAGCTCTTCGCCT TTACGCATGCCCTGAGCCAGAGC TACGGTGCAGGAGAGC | *P. inhibens* P72 | 57 kb plasmid reporter strain | Current study |
| 132 | CGTAGCACCAGGCGTTTAAGG | pBBR1MCS-5 plasmid | Validation of reporter strains | Current study |
| 255 | ACTTGAAGAAGTCATGCTGC | sfGFP protein | Validation of reporter strains | Current study |

| | | | | |
|---|---|---|---|---|
| 1169 | GTACAAAAAAGCAGGCTCCGAATTCGCCCTTAACCAGTCCTTCCGCAACAAGACGCGCAAGG | *P. inhibens* DSM 17395 | Construction of the 65 kb plasmid mutant | Current study |
| 1170 | CAGTTTACGAACCGAACAGGCTTATGTCAAGCGAAATATCACTATCTCCACACTACCACC | *P. inhibens* DSM 17395 | Construction of the 65 kb plasmid mutant | Current study |
| 1171 | CTTCTATCGCCTTCTTGACGAGTTCTTCTGAGAGGATTAATTTAAATCGCACTCTTATTCTC | *P. inhibens* DSM 17395 | Construction of the 65 kb plasmid mutant | Current study |
| 1172 | CTTTGTACAAGAAAGCTGGGTCGAATTCGCCCTACCATTCATAAGCACGTGCTGTGGAAATGTC | *P. inhibens* DSM 17395 | Construction of the 65 kb plasmid mutant | Current study |

**Table S5:** Primer list of bacterial genes for qRT-PCR.

| Strain | Genetic locus | Gene name | Accession number | Forward Primer | Reverse Primer |
|---|---|---|---|---|---|
| *D. shibae* DFL-12 | Chromosome | *recA* | DSHI_RS08365 | CCGTAATCTCGA GAGCCTGC | GAAAGTTCCGG CAAGACCAC |
| | Chromosome | *gyrA* | DSHI_RS07515 | GTGAAGCGGTA GAGCTGGTTc | CTCGACGAGAT CCCCTACCA |
| | 191 kb and 126 kb plasmids | *virB8* | DSHI_RS18440 and DSHI_RS20035 | TTCCGCTATGTG ACCGACC | CGATCCGGTTG ATGCTGAG |
| | 191 kb and 126 kb plasmids | *virB4* | DSHI_RS18425 and DSHI_RS20020 | CGATCTGACCG GCATTCTC | GTCACCAGCCA GTTCGACAG |
| | 191 kb and 126 kb plasmids | *virD4* | DSHI_RS18490 and DSHI_RS20080 | GAGCATGATCAT CACGGGc | CGGTGTCGGA CTTTGATTTC |
| *P. inhibens P72* | Chromosome | *recA* | PhaeoP72_RS07860 | GCTGACACCCA AGTCGGAG | AGCCGAACATA ACGCCAATCT |
| | Chromosome | *gyrA* | PhaeoP72_RS07800 | TCAGCTTATGTC GGGCTTCG | CGGTTCCTGAC CTCCTTCC |
| | 18 kb and 57 kb plasmids | *virB8* | PhaeoP72_RS19740 and PhaeoP72_RS20905 | GGGTTCGTCCA GACCTCATC | CCGGCTCGAT GTGGAAAT |
| | 18 kb and 57 kb plasmids | *virB4* | PhaeoP72_RS19755 and PhaeoP72_RS20885 | ACCGGCATTCT CGACAGC | GCGTATTGCGT CATCATCAC |
| | 18 kb and 57 kb plasmids | *virD4* | PhaeoP72_03985 and PhaeoP72_RS20940 | GCTTCGACCAAT AACGACACG | GAAGAGGCGG ATCAGCGG |

**Table S6**: Plasmids used in the study.

| Plasmid name | Origin | Comments |
|---|---|---|
| pBBR1MCS-5 | Prof. Kenneth M. Peterson, Louisiana State University Medical Center, USA[82] | |
| pYDR1 | Lab collection | Constructed using pBBR1MCS-5 and used in the current study as a vector carrying the sfGFP protein |
| pYDR3 | Current study | Constructed using pYDR1 by replacing the sfGFP promoter with the 184 kb plasmid *virB* promoter for *P. inhibens* P72 reporter strain |
| pYDR10 | Current study | Constructed using pYDR1 by replacing the sfGFP promoter with the 57 kb plasmid *virB* promoter for *P. inhibens* P72 reporter strain |
| pYDR6 | Current study | Constructed using pYDR1 by replacing the sfGFP promoter with the 126 kb plasmid *virB* promoter for *D. shibae* DFL-12 reporter strain |
| pYDR7 | Current study | Constructed using pYDR1 by replacing the sfGFP promoter with the 191 kb plasmid *virB* promoter for *D. shibae* DFL-12 reporter strain |
| pCR™8/GW/TOPO | Invitrogen | Cat#K250020 |
| pYDR8 | Lab collection | Constructed using pCR8/GW/TOPO vector (Invitrogen), and used in the current study as a vector carrying the kanamycin resistant cassette |
| pYDR9 | Current study | Constructed using pYDR8 vector for creating the 65 kb intergenic mutant of *P. inhibens* DSM 17395 |

**Table S8. Quantification of donors, recipients, and transconjugants.** Each conjugation donor-recipient pair from the conjugation assay described in figure 5 was plated on selective plates in serial dilutions, and quantified. The numbers indicate the colony-forming units per milliliter (CFU/ml). CFU values represent an average of two technical replicates for each biological sample for the donors and transconjugants (for conjugation efficiency calculations), and one technical replicate for the recipients. Dilution- indicates the dilution factor that was plated on the selective plates. ND- not detected and omitted from the average. Donor plasmid denoted as "p" is indicated by its size (Kb), recipients denoted as "chr", antibiotic resistance markers are indicated by gent for gentamicin-resistance and kana for kanamycin-resistance.

| Donor strain | Recipient strain | Treatment | Biological replicate | Donor CFUs/ml | Dilution | Recipient CFUs/ml | Dilution | Transconjugants CFUs/ml |
|---|---|---|---|---|---|---|---|---|
| *D. shibae* DFL-12 $^{chr-gent}$ | *P. inhibens* DSM 17395$^{chr-kana}$ | Control | 1 | 6.83E+08 | -5 | 2.50E+07 | -4 | 4.55E+03 |
| | | | 2 | 7.70E+08 | -5 | 3.20E+07 | -4 | 4.30E+03 |
| | | | 3 | 7.65E+08 | -5 | 2.00E+07 | -4 | 7.53E+03 |
| | | | 4 | 6.10E+08 | -5 | 2.40E+07 | -4 | 3.05E+03 |
| | | Algal filtrates | 1 | 5.70E+08 | -5 | 4.20E+07 | -4 | 8.40E+03 |
| | | | 2 | 6.40E+08 | -5 | 2.70E+07 | -4 | 5.90E+03 |
| | | | 3 | 5.65E+08 | -5 | 4.40E+07 | -4 | 7.75E+03 |
| | | | 4 | 6.60E+08 | -5 | 3.30E+07 | -4 | 8.80E+03 |
| *P. inhibens* P72 $^{p57-kana}$ | *D. shibae* DFL-12 $^{chr-gent}$ | Control | 1 | 6.00E+07 | -4 | 2.90E+08 | -5 | 6.35E+03 |
| | | | 2 | 5.75E+07 | -4 | 4.10E+08 | -5 | 4.75E+03 |
| | | | 3 | 5.90E+07 | -4 | 2.80E+08 | -5 | 4.80E+03 |
| | | | 4 | 6.70E+07 | -4 | 2.60E+08 | -5 | 4.55E+03 |
| | | Algal filtrates | 1 | 1.00E+08 | -4 | 5.40E+08 | -5 | 9.65E+03 |
| | | | 2 | 1.50E+08 | -4 | 6.70E+08 | -5 | 8.30E+03 |
| | | | 3 | 9.13E+07 | -4 | 4.80E+08 | -5 | 9.00E+03 |
| | | | 4 | 9.33E+07 | -4 | 5.25E+08 | -5 | 2.25E+04 |
| *P. inhibens* P72 $^{p57-kana}$ | *M. algicola* DG898 $^{chr-gent}$ | Control | 1 | 6.65E+07 | -4 | 4.90E+08 | -5 | 2.05E+03 |
| | | | 2 | 7.50E+07 | -4 | ND | -5 | 1.20E+03 |
| | | | 3 | 7.85E+07 | -4 | 4.50E+08 | -5 | 1.45E+03 |
| | | | 4 | 7.35E+07 | -4 | 4.80E+08 | -5 | 3.05E+03 |
| | | Algal filtrates | 1 | 9.05E+07 | -4 | 3.00E+08 | -5 | 1.06E+04 |
| | | | 2 | 7.95E+07 | -4 | 3.60E+08 | -5 | 1.14E+04 |
| | | | 3 | 8.60E+07 | -4 | 2.80E+08 | -5 | 1.35E+04 |
| | | | 4 | 8.05E+07 | -4 | 4.60E+08 | -5 | 2.08E+04 |
| *P. inhibens* P72 $^{p57-kana}$ | *P. inhibens* DSM 17395$^{chr-gent}$ | Control | 1 | 1.20E+08 | -4 | 1.20E+08 | -5 | 1.38E+05 |
| | | | 2 | 9.00E+07 | -5 | 7.00E+07 | -5 | 1.02E+05 |
| | | | 3 | 9.00E+07 | -4 | 1.00E+08 | -5 | 1.40E+05 |
| | | | 4 | 9.00E+07 | -5 | 1.30E+08 | -5 | 2.50E+05 |
| | | Algal filtrates | 1 | 1.00E+08 | -4 | 1.60E+08 | -5 | 3.45E+05 |
| | | | 2 | 1.60E+08 | -5 | 1.20E+08 | -5 | 5.65E+05 |
| | | | 3 | 1.60E+08 | -5 | 1.40E+08 | -5 | 4.40E+05 |
| | | | 4 | 2.10E+08 | -5 | 1.50E+08 | -5 | 6.00E+05 |
